# The Origin and Evolution of Sex Peptide and Sex Peptide Receptor Interactions

**DOI:** 10.1101/2023.11.19.567744

**Authors:** Junhui Peng, Nicolas Svetec, Henrik Molina, Li Zhao

**Affiliations:** Laboratory of Evolutionary Genetics and Genomics, The Rockefeller University, New York, NY, USA; Proteomics Resource Center, The Rockefeller University, New York, NY 10065, USA

## Abstract

Post-mating responses play a vital role in successful reproduction across diverse species. In fruit flies, sex peptide (SP) binds to the sex peptide receptor (SPR), triggering a series of post-mating responses. However, the origin of SPR predates the emergence of SP. The evolutionary origins of the interactions between SP and SPR and the mechanisms by which they interact remain enigmatic. In this study, we used ancestral sequence reconstruction, AlphaFold2 predictions, and molecular dynamics simulations to study SP-SPR interactions and their origination. Using AlphaFold2 and long-time molecular dynamics (MD) simulations, we predicted the structure and dynamics of SP-SPR interactions. We show that SP potentially binds to the ancestral states of Diptera SPR. Notably, we found that only a few amino acid changes in SPR are sufficient for the formation of SP-SPR interactions. Ancestral sequence reconstruction and MD simulations further reveal that SPR interacts with SP through residues that are mostly involved in the interaction interface of an ancestral ligand, myoinhibitory peptides (MIPs). We propose a potential mechanism whereby SP-SPR interactions arise from the pre-existing MIP-SPR interface as well as early chance events both inside and outside the pre-existing interface that created novel SP-specific SP-SPR interactions. Our findings provide new insights into the origin and evolution of SP-SPR interactions and their relationship with MIP-SPR interactions.

## Introduction

Reproductive success plays a crucial role for individuals, populations, and species. Predominantly, males of many species undergo intense sexual selection because of their necessity to secure mating partners successfully, and higher intrasexual variance in mating success among males. Inter-male competition emerges as a potent force propelling sexual selection. This competition is manifested at the phenotypic level through distinct male attributes such as large horns in beetles or vivid colors in birds (Eberhard, 1985; Hopkins & Kopp, 2021; Parker, 1970). Moreover, at the molecular level, this competitive landscape stimulates the accelerated evolution of genes, which may subsequently contribute to the development of phenotypes that either precipitate or resolve sexual conflict (Ferris et al., 1997; Haerty et al., 2007; Kokko & Jennions, 2014), where genotypical traits beneficial by one sex might be harmful to the other (Chapman et al., 2003).

In many insects, the mating behavior of females, including their potential to mate again, affects the evolution of male reproductive and genotypical traits. Males aim to increase their chance of mating and sperm numbers or competitivity, and if possible, prevent their female partners from mating again. For example, in fruit flies, males have evolved molecular mechanisms that cause changes in females that increase male reproductive success under competitive mating conditions. This is achieved by transferring seminal fluid proteins (SFPs) and other components to the female during mating. Within mated females, SFPs can induce a range of crucial post-mating responses and functions, such as increasing egg production, stimulating the immune system, and reducing the female’s sexual receptivity (Chapman, 2001; Clark et al., 1995; Findlay et al., 2014; Haerty et al., 2007; Ram & Wolfner, 2007, 2009; Sirot et al., 2015; Wigby et al., 2020; Wolfner, 1997, 2002). These seminal fluid proteins are often rapidly evolving and undergo substantial adaptive changes within a species (Begun et al., 2000; Begun & Lindfors, 2005; Findlay et al., 2008, 2009; Kern et al., 2004; Wagstaff & Begun, 2005).

In the fruit fly, *D. melanogaster*, the sex peptide (SP) is a seminal fluid protein that plays a pivotal role in initiating post-mating responses in mated females (Chen et al., 1988; Hopkins & Perry, 2022). After cleavage of the 19 amino-acid N-terminal signal peptide in the precursor, mature SP is a small protein with 36 amino acids. Despite its small size, it possesses multiple functional domains. In the mature 36 amino-acid peptide, the N-terminal part is responsible for juvenile hormone induction and for binding to sperm (Peng, Chen, et al., 2005). The proline-rich motif in the middle part is necessary for stimulating the immune system (Domanitskaya et al., 2007; Peng, Zipperlen, et al., 2005). The C-terminal end enables changes in oviposition and receptivity by binding to the sex peptide receptor (SPR) and further interacting with neuronal circuits through sensing neurons (SPSNs) (Feng et al., 2014; Rezával et al., 2012). In addition to enhancing oviposition and reducing the receptivity of females (Aigaki et al., 1991; Avila et al., 2010; Chapman, Bangham, et al., 2003; Chen et al., 1988; Liu & Kubli, 2003), SP can induce numerous other behavioral and physiological changes (Chen et al., 1988; Hopkins & Perry, 2022; Wigby & Chapman, 2005), such as enhancing long time memory (Scheunemann et al., 2019), stimulating immune responses (Peng, Zipperlen, et al., 2005), affecting midgut morphology and physiology (White et al., 2021), and regulating circadian rhythms (Delbare et al., 2023) in females.

Despite SP’s fundamental role in these post-mating scenarios, its evolutionary origins and the intricate mechanisms governing its interaction with SPR remain enigmatic. Adding to this complexity is that the existence of SPR predates the emergence of SP (Kim et al., 2010; Poels et al., 2010; Yapici et al., 2008), which raises further questions about the origin of SP and its interactions with SPR. On the other hand, SPR, a gene conserved among various *Insecta* species, such as in mosquitoes, beetles, and moths, is known to bind with conserved myoinhibitory peptides (MIPs) (Kim et al., 2010; Poels et al., 2010). MIPs are small peptides acting as neuromodulators that inhibit muscle activity while simultaneously serving roles in several other physiological processes (Oh et al., 2014).

The transition from a singular MIPs-SPR interaction to a dual MIPs-SPR and SP-SPR interaction in *Drosophila* poses a significant unanswered question. To address these queries, we utilized ancestral sequence reconstruction, AlphaFold2 multimer predictions (Evans et al., 2022; Jumper et al., 2021), and molecular dynamics (MD) simulations to investigate the biophysical interactions between SP and SPR, and to trace their evolutionary origins. Our objective was to elucidate the complex dynamics and evolutionary trajectory of SP and SPR interactions, thereby enhancing our understanding of post-mating responses and the underlying molecular and biochemical alterations.

## Result

### C terminus of SP binds stably in the SPR extracellular pocket from AlphaFold2 multimer predictions and MD simulations

SP has been shown to bind to SPR through its C-terminus (Kim et al., 2010; Schmidt et al., 1993; Yapici et al., 2008). However, the detailed molecular mechanism was unclear. Here, we used AlphaFold2 to predict the binding mode of SP to SPR. From our predictions, SP binds to SPR with relatively high confidence, with interface pTM (ipTM) of 55, predicted aligned error (PAE) of 2.3 Å (Figure S1), and pDockQ of 0.47. A pDockQ value exceeding 0.23 and close to 0.5 indicates that our SP-SPR complex model is highly acceptable (Bryant et al., 2022; Burke et al., 2023). In the predicted SP/SPR model, the C terminus of SP (designated as SP_17-36 in Figure 1, corresponding to SP residues 17-36) inserted into the extracellular ligand-binding pocket of SPR (Figure 1A, 1B, Fig. S1A, S1B), which is consistent with earlier biochemical and molecular biology studies (Kim et al., 2010; Schmidt et al., 1993; Yapici et al., 2008). To further evaluate the stability and dynamics of SP/SPR interactions, we conducted three independent MD simulations totaling 14 μs starting from the predicted SP_17-36/SPR complex (Fig S1C, *Methods*). Throughout the MD simulations, SP remained tightly bound to the SPR extracellular pocket (Fig. 1C, Fig. S1D), demonstrating that SP_17-36/SPR is overall stable. We computed the root mean square deviation (RMSD) of SP after aligning the trajectory using SPR transmembrane domains. The RMSD values of SP_17-36 ranged from 2 to 6 Å, indicating some degree of flexibility of SP in the bound state (Figure 1C, Fig. S1D). Intriguingly, when we further computed the RMSD of SP_22-36, we observed that this core region was extremely constrained and rigid with RMSD around 2 Å (Figure 1C, Fig S1D). This suggests that the stable SP/SPR interactions were mostly regulated by the most C-terminus 15 residues, which we term the C-terminal core regions. This finding is in line with earlier biochemical experiments (Ding et al., 2003; Kim et al., 2010; Schmidt et al., 1993). Upon further detailed structural analysis, we found about 24 residues in SPR frequently interacted with the SP core regions at high frequencies (>50%, Figure 1D, *Methods*) in all the three independent MD simulations. These interactions include salt-bridge, pi-cation, aromatic ring stacking, and hydrophobic interactions. The salt-bridge interactions include D82, E172, and D266 of SPR with R35, R35, and K22 of SP, respectively, while the pi-cation interactions include R276 and H352 of SPR with W23 and W32 of SP, respectively. Notably, several residues interact with more than one SP residues (Figure 1D and 1E). For example, Y273 of SPR interacts with both W23 and W32 in SP (Figure 1D-E) through aromatic ring stacking interactions, and the two W residues in SP were previously suggested to be important for SP-SPR interactions (Kim et al., 2010). Besides the two tryptophan residues, Y273 also interacts with G35, R35, and C36 at high frequencies (Figure 1D-E) through hydrophobic interactions. Another example is H352, which frequently interacts with residue W23 of SP through pi-cation interactions and residues K22 and L26 of SP through hydrophobic interactions. Collectively, these extensive interactions enable SP to bind tightly to SPR.

**Figure 1.**
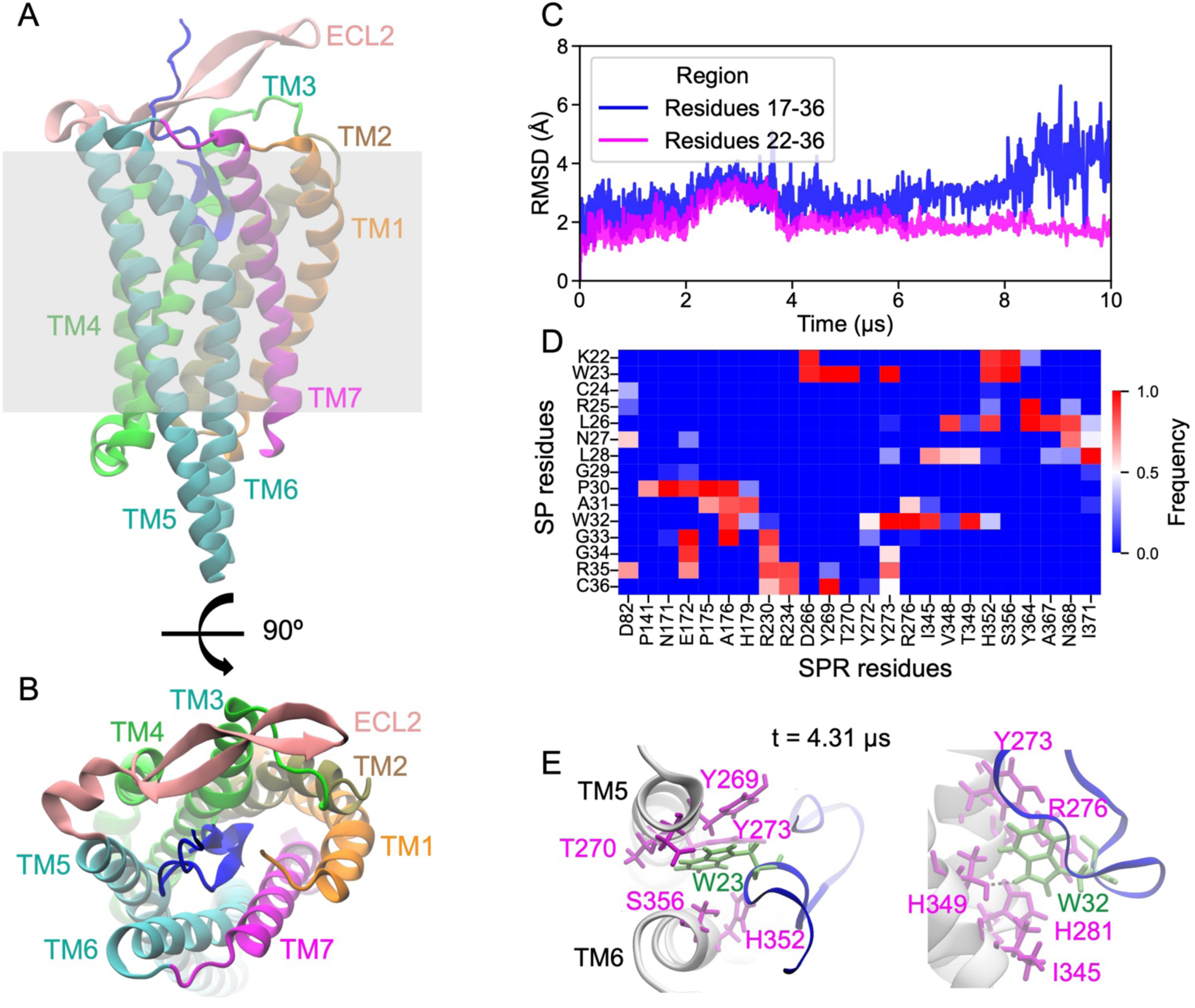
Structure and dynamics of SP/SPR complex. (A) Structural model of SP/SPR complex predicted by AlphaFold2, where SP is colored in blue and SPR is colored in different colors according to the transmembrane segments, starting from TM1 to TM7. The extracellular loop 2 (ECL2) of SPR is also highlighted. (B) Top view of (A). (C) Time resolved RMSD of the C-terminus (blue) and core regions (magenta) during MD simulations. The small RMSD values of the core regions suggest a stable SP/SPR interaction. (D) Frequently interacting residues between SP (residues labeled on Y axis) and SPR (residues labeled on X axis) during MD simulations (frequency > 50% in all three independent MD simulations). (E) A representative structure at t = 4.31 μs from the 10-μs MD simulation, showing the interactions between SPR residues (magenta) with W23 and W32 in SP (green).

### Sex peptide genes are widespread in *Drosophila* genus and older than previously estimated

The *SP* gene, which codes for sex peptide (SP), has been reported for decades (Chen et al., 1988). It was later reported to be specific to a subset of *Drosophila* species (Kim et al., 2010; Tsuda & Aigaki, 2016; Yapici et al., 2008). To gain a deeper understanding of the origin of the *SP* gene, we took advantage of the conservation of the C-terminal sequences of SP. We applied *tblastn* (Altschul et al., 1990) and gene-structure aware alignments (Iwata & Gotoh, 2012) to search against NCBI eukaryotes representative genomes (Sayers et al., 2022) (Material and Methods). We identified 61 *SP* genes and their duplications in 33 representative genomes (Figure 2A, supplementary file S1). Our results show that the *SP* gene can be found mostly in *Drosophila* species and date its origin to at least the common ancestor with *Scaptodrosophila lebanonensis*. This suggests that the *SP* gene age is much older than previously reported (Tsuda & Aigaki, 2016). Our findings are consistent with a recent study by Hopkins et al. (Hopkins et al., 2024), who also identified one copy of *SP* gene in *S. lebanonensis.* Interestingly, by analyzing RNA-seq data (https://www.ncbi.nlm.nih.gov/bioproject/PRJNA554780), we found that the *SP* gene in *S. lebanonensis* is expressed specifically in male gonads (Figure S2). Although it is unclear about the function of SP in *S. lebanonensis* (Tsuda et al., 2015; Tsuda & Aigaki, 2016), the apparent gonad-specific expression suggests that it may function partially similarly to *D. melanogaster* SP.

**Figure 2.**
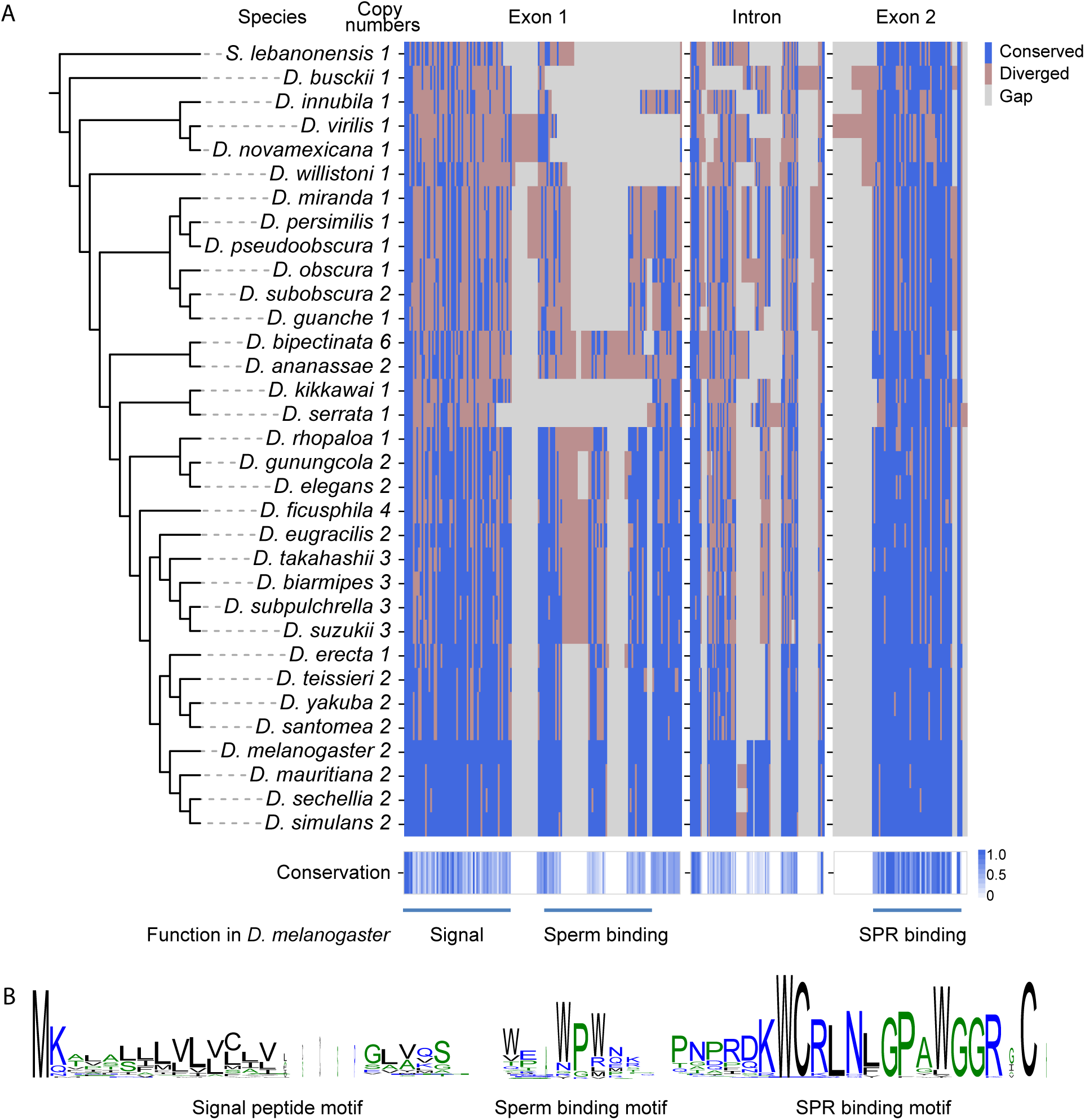
Sequence conservation of *SP* genes. (A) Phylogenetic tree of the species that have *SP* genes identified. The tree was extracted from timetree.org (Kumar et al., 2022). The left panel shows the phylogenetic tree of the 33 species in NCBI representative genomes with *SP* genes identified. The copy number of *SP* genes in each species is listed after the species name. The right panel shows the alignment of the two exons and intron of the representative *SP* genes. For species that have more than one copy of *SP* genes, the representative *SP* gene in each species has the highest sequence similarity to the *SP* gene in *D. melanogaster*. Alignment gaps were shown in gray, while diverged and conserved nucleotides were shown in brown and blue, respectively. The bottom panel shows the conservation of the alignments, where the functional regions that code for signal peptide, sperm binding sites, and SPR binding sites in *D. melanogaster* are highlighted. (B) Sequence of the SP proteins across the 33 species. The N-terminal signal peptides were enriched in hydrophobic residues (left panel). The putative sperm binding motif in *D. melanogaster* is WEWPWNR, while it can be quite different in other species (middle panel). The SPR binding motif is highly conserved with two tryptophan residues (right panel).

Across the identified species, the *SP* genes preserve similar gene structures, each with two exons and one short intron. The length of the short intron ranges from 51 bp to 75 bp. Interestingly, while the first exon is diverged, all *SP* genes are predicted to harbor a 5’ region coding for an N-terminal signal peptide (Figure 2B, Figure S3). The region coding for the sperm binding motif (Peng, Chen, et al., 2005) *in D. melanogaster* follows the N-terminal signal peptide and is highly diverged in *Drosophila* subgenus species (Figure 2B, Figure S3). This might suggest that SP in these species: (1) may not have evolved the ability to bind to sperm; or (2) may bind to sperm through different mechanisms as reported in *D. melanogaster*, where a different set of protein interactions could be involved (Singh et al., 2018). The second exon is short and highly conserved, and the peptide products were reported to be responsible for SPR binding in *D. melanogaster* (Kim et al., 2010). The highly conserved sequences suggest that SP from other species may have the capability to bind to SPR, provided that SPR is expressed in the female reproductive tract or both SP and SPR are expressed in the same tissue (Figure 2B, Figure S3). A similar conservation pattern has been observed in a set of genome assemblies from ∼145 *Drosophila* species in a recent study (Hopkins et al., 2024).

Most species within the *Drosophila* subgenus have only one copy of *SP* gene, whereas most species in *Sophophora* subgenus contain two or more copies, suggesting a recent expansion of SP genes in these species. Some species in *Sophophora* subgenus even have three or more copies. For example, *D. bipectinata* in the *ananassae* subgroup have 6 copies, and most of the *suzukii* subgroup species contain three copies. The results suggest frequent gene copy number variation events in *Sophophora* subgenus species.

To gain a deeper insight into the origins of *SP* gene copy number variations, we constructed the phylogenetic tree of all the identified 61 *SP* genes (Figure S4). The phylogenetic tree revealed that *Dup99B gene* copy in *D. melanogaster* could date back to anannassae subgroup species, as all *SP* genes in anannassae subgroup (2 in *D. anannassae* and 6 in *D. bipectinata*) were closer to *Dup99B* than *SP* copy in *D. melanogaster*. Our results also suggested a complex evolutionary history of *SP* genes, where Dup99B copy might be lost in *D. elegans* and SP copy might experience multiple duplication events in suzukii subgroup, gunungcola, and some other *Drosophila* species (Figure S4). Furthermore, we found that two of the three copies in suzukii subgroup species are recent suzukii subgroup specific tandem duplications, whereas the third copy in suzukii subgroup species shares a similar origin as *Dup99B*. Interestingly, we found that the high numbers of copies in *D. bipectinata* were anannassae subgroup specific. These results suggest that *SP* genes were likely to experience a substantial expansion in the anannassae subgroup species. Many of these high copy numbers might be lost due to weaker selective constraints. Two copies were maintained in melanogaster group species, i.e., the *SP* gene copy and the *Dup99B* gene copy. Among the two copies, the *Dup99B* copies were highly diverged while the *SP* copies were highly conserved and were further duplicated in the suzukii subgroup.

Within the suzukii species complex, three copies of the sex peptides are present. To investigate whether they are translated, we conducted mass spectrometry and targeted mass spectrometry on accessory gland samples from *D. suzukii* and *D. subpublchrella*. For *D. suzukii*, we identified one unique peptide corresponding to XP_016924330.1, two unique peptides for XP_016924329.1, and two peptides shared between XP_016924329.1 and XP_016924330.1 (Figure S5). In the case of *D. subpubchrella*, four peptides matched both XP_037720360.1 and XP_037721024.1 (Figure S5). These findings indicate that duplicated copies of sex peptides are translated, either at similar levels or with one copy being more highly transcribed or translated than the other. Future research could intriguingly explore the potential functional divergence of these genes.

### MIP-SPR share a similar binding interface with SP-SPR

MIPs have been identified as ancestral ligands for SPR (Kim et al., 2010). The MIP precursor can be cleaved into five MIP peptides, MIP1, MIP2, MIP3, MIP4, and MIP5. Here, we used AlphaFold2 to predict the structures of MIP-SPR complexes and ran MD simulations to study the dynamics of the interactions. To compare the SPR interface for SP and MIP peptides, we extracted residues in SPR that interact with SP and MIP peptides at a frequency greater than 5%, respectively. Interestingly, we found that all five MIP-SPR complexes share similar binding interfaces to the SP-SPR complex (Figure 3). In general, SP shares many interacting sites in SPR with MIP peptides (Figure 3A and 3B). For example, D82, E172, R230, R234, L253, T255, V265, D266, Y269, T270, Y273, H352, S356, Y364, N368, and I371 interact with SP and all the five MIPs (Figure 3C). Many of the residues are shared by SP and at least one MIP peptide, such as W77, L79, S83, Y92, T137, P141, L145, N171, P175, A176, H179, I345, V348, T349, I353, L357, E360, L362, A367, M372, and N375 (Figure 3C). By examining SP-interacting residues in SPR during the MD simulations, we pinpointed some SPR residues that interact with SP (frequency > 5%), with no or very low-frequency interactions with MIP peptides. These SPR residues include M69, E70, N78, T173, Q227, D233, Y272, R276, H281 (Figure 3C). Although MIP-SPR and SP-SPR interactions are mostly similar, the fact that SP and MIP potentially have their unique interface was supported by a recent study examining ligand selectivity against SPR in *D. melanogaster* and *Aedes aegypti*, where the authors found a few amino acids can be important for SP and MIP sensitivity, respectively (J.-H. Lee et al., 2020). We extracted SPR residues that frequently interacts with those two residues to investigate whether the two conserved tryptophan residues of the W(X)_n_W motifs in SP and MIP peptides share similar interaction patterns. Interestingly, we found that the first tryptophan residue in SP and all five MIP peptides showed similar interaction patterns (Figure S6). For the second tryptophan residue, all the five MIP peptides share similar interaction patterns, while SP displayed a different pattern but shared some interacting residues in SPR (Figure S6). Overall, our findings further support that the origin of SP-SPR interactions might arise from a combination of exploiting pre-existing MIP-SPR interfaces and the evolution of new SP-SPR interactions.

**Figure 3.**
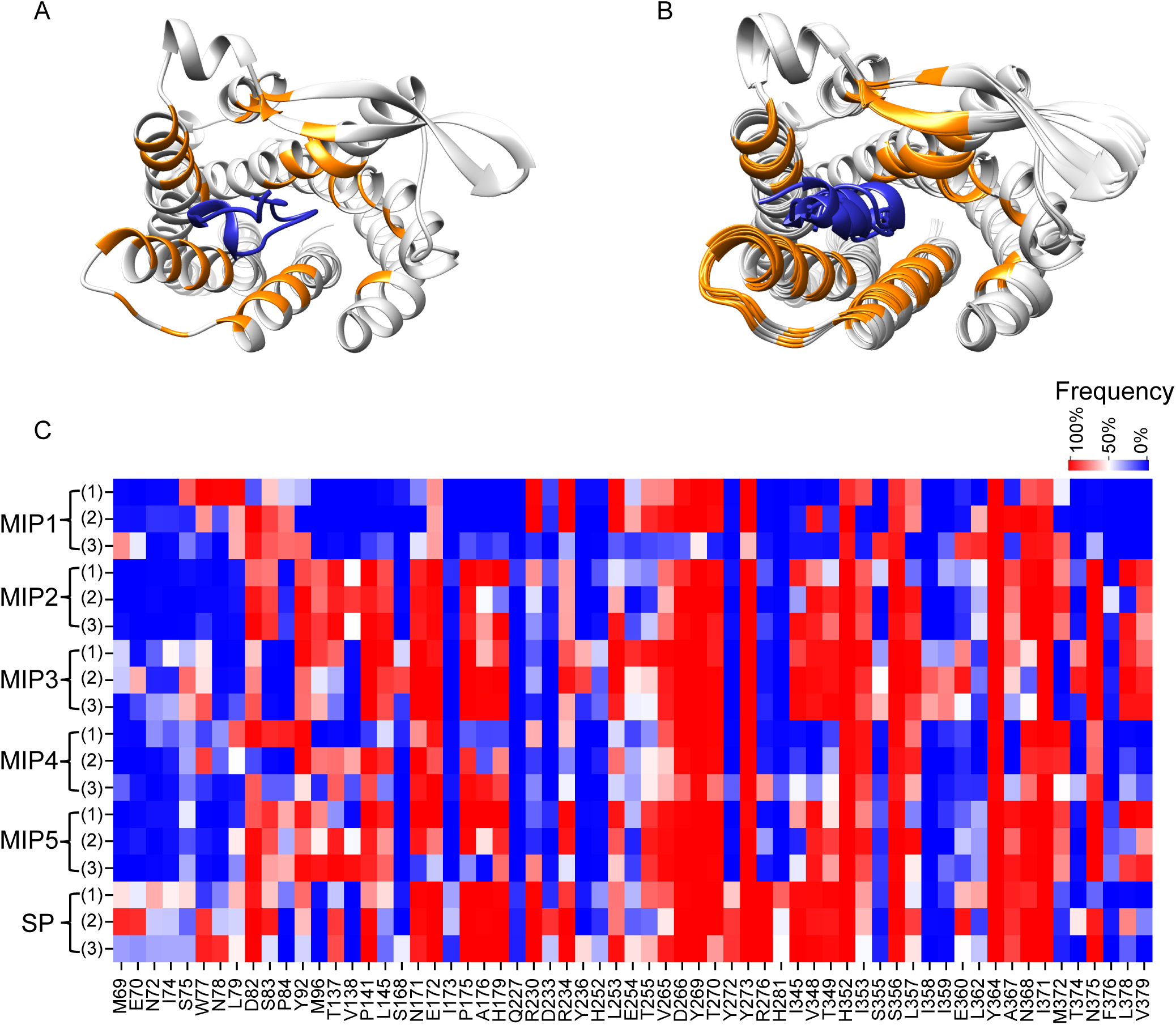
Mip/SPR shares a similar binding interface with SP/SPR. (A) Top view of SP-SPR complex. SP is colored in blue and the SPR residues that interact with SP in MD simulations are colored in orange. (B) Top view of the five Mip-SPR complexes overlaid together. Mip peptides are colored in blue and the SPR residues that interact with SP in MD simulations are colored in orange as in (A). (C) The frequency of SPR residues interacting with SP or MIPs during MD simulations. SPR residues that interact with either SP or MIPs at 5% frequency or higher during MD simulations were shown.

### A few pre-*Drosophila* SPR mutations might create favorable conditions for the origination of SP/SPR interactions

We collected and aligned SPR sequences across *Metazoa* species from orthoDB (Kuznetsov et al., 2023) (Material and Methods). The SPR sequences showed a highly diverged N-terminus and relatively conserved core regions (Figure S7). We then extracted the residues in SPR that frequently interact with SP and MIP during MD simulations (see section *MIP-SPR share a similar binding interface with SP-SPR* in Results). We found that most of these residues were highly conserved in the 38 *Drosophila* species examined in our study, suggesting strong selective constraints in SPR against SP binding so that the predicted SP-interacting residues in SPR were constrained and highly conserved. The number of conserved amino acids decreased sequentially in *Diptera*, *Insecta*, *Arthropoda*, and *Metazoa* species (Figure 4A), suggesting that these sites might undergo decreased selective constraints.

**Figure 4.**
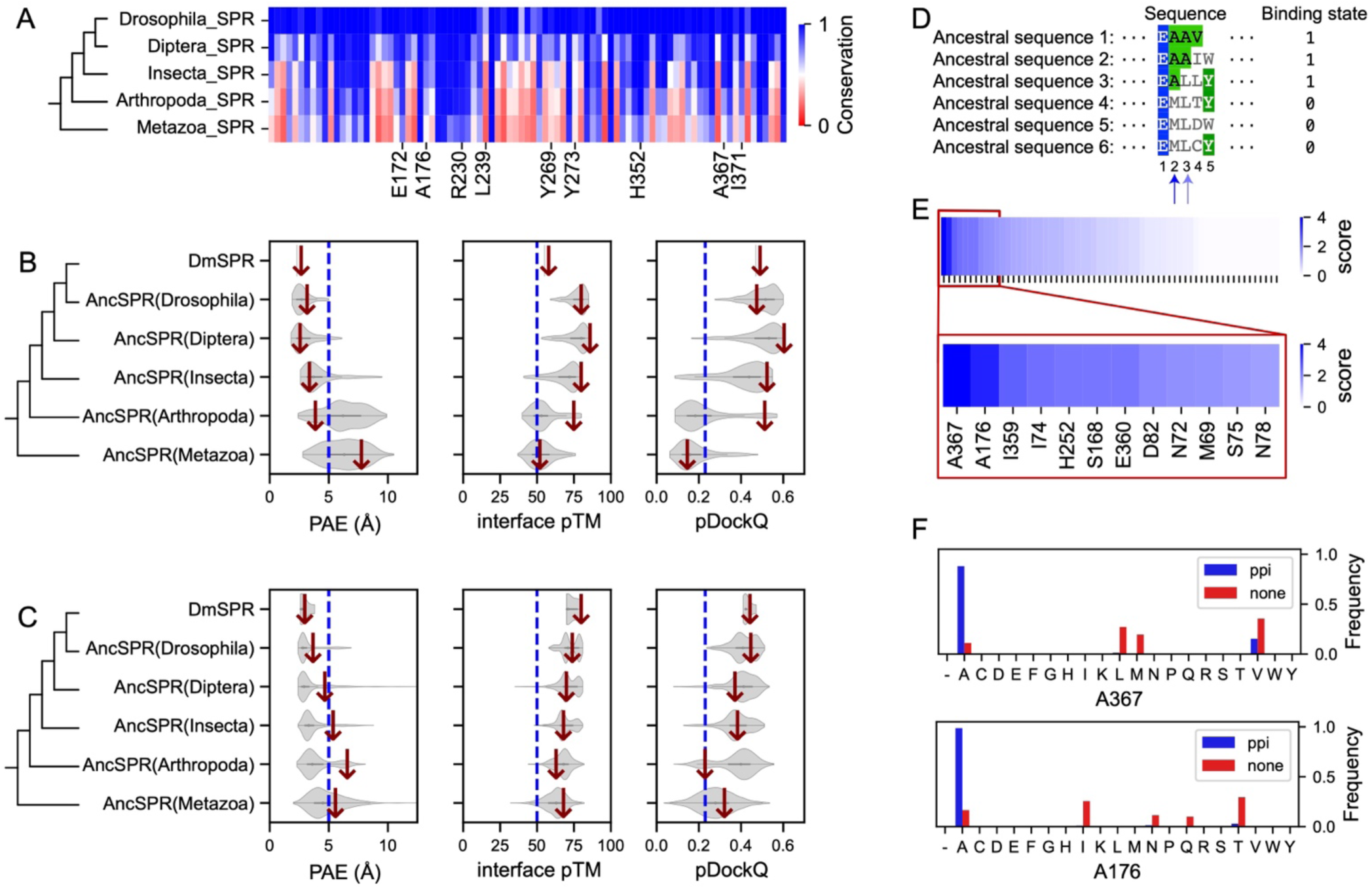
In silico coevolution between SPR residues and SP binding states. (A) Conservation of SP and Mip binding sites in SPR. (B) AlphaFold predictions of SP and ancestral SPRs binding. (C) AlphaFold predictions of Mips and ancestral SPRs binding. (D) Simplified scheme of the *in-silico* coevolution analysis. We computed whether there are sites that co-varied with SP binding states. Binding state of 1 suggests acceptable binding potential between the ligand and receptor, 0 for not binding. In this simplified example, site 2 has a high chance to co-vary with binding state, followed by site 3, but not sites 1, 4, and 5. (E) Co-varying scores of SPR residues and SP binding states. A367 and A176 showed the highest score. (F) The frequency of each amino acid at residue sites 367 and 176 in different SP binding states.

To find out which changes in SPR from old lineages to *Drosophila* lineage might result in SP-SPR interactions, we first reconstructed the ancestral states of SPR of *Drosophila*, *Diptera*, *Insecta*, *Arthropoda*, and *Metazoa* (Material and Methods). For each of the five ancestors, we extracted the most probable ancestral sequences and randomly sampled another one hundred alternative ancestral sequences, respectively (Material and Methods). We used AlphaFold2 to predict whether the ancestral SPRs could bind to SP or MIPs. Interestingly, we found that most of the ancestral *Insecta* SPRs could potentially bind to SP (pDockQ greater than 0.23), and the SP binding potential decreased drastically for ancestral *Arthropoda* and *Metazoa* SPRs (Figure 4B). In contrast, a similar trend was not observed in the case of MIP peptides binding potentials (Figure 4C).

We then computed whether these predicted SP-interacting sites in SPR showed high co-varying scores with their SP binding states (illustrated in Figure 4D) by an in-silico co-varying approach (Material and Methods). We found that several residues showed high co-varying scores (Figure 4E), suggesting that these sites co-varied with SP binding states at high probabilities. Among these residues, A367 and A176 showed the highest scores. In the SP binding states, the two sites were both Alanine. However, in the none-SP binding states, the two sites are more likely to be other amino acids, for example, Leu, Met, and Val at site 367, and Ile and Thr at site 176 (Figure 4F). We then used AlphaFold2 to predict whether the top-scored residues could affect SP-SPR binding. Interestingly, with only the top scored six amino acids changed to their alternative ancestral states, the mutated sequence did not show SP binding potential, with pDockQ lower than 0.23 (Figure S8). A pDockQ value of 0.23 is the acceptable pDockQ cutoff to define interaction (Bryant et al., 2022). These results indicate the co-varied residues alone could potentially switch on/off SP-SPR binding.

Since A367 and A176 were in the SP binding interface (Figure 5A), we analyzed the interactions between A367 and A176 in SPR with each of the SP residues during the 10-μs MD simulations to further check whether the two sites could affect SP-SPR interactions (Material and Methods). We found that A367 showed frequent interactions with L26 and L28 with the minimum distances frequently ranging between 6 Å and 3.5 Å (Figure 1D, Figure 5B). A distance greater than 6 Å suggests a lack of interaction, while a distance below 3.5 Å is often indicative of polar interactions. Notably, distances of nonpolar residues that lie between 6 Å and 3.5 Å are characteristic of hydrophobic interactions (Material and Methods). Meanwhile, A176 showed frequent interactions with P30, A31, W32, and G33 (Figure 1D, Figure 5C). The results suggest that A367 and A176 form a binding pocket for SP and closely interact with SP through hydrophobic interactions (Material and Methods). Mutations of these two residues from Alanine to other residues, e.g., Leucine, Methionine as shown in Figure 4F, could potentially introduce steric hindrance between SP and SPR due to their large side chains compared to Alanine. The steric hindrance could further reduce the SP binding potentials for SPR harboring those mutations. Interestingly, in the ancestral states of *Arthropoda* and *Metazoa* SPR, where most of them did not show SP binding potential, the two sites were mostly not Alanine (Figure 5D, Figure S9). In *Drosophila* the two sites were both Alanine, and in *Diptera* and *Insecta*, the two sites were highly conserved and were mostly Alanine (Figure 5D). The results further indicate that the pre-Drosophila changes of these residues to Alanine could potentially create changes for the origination of SP-SPR interactions.

**Figure 5.**
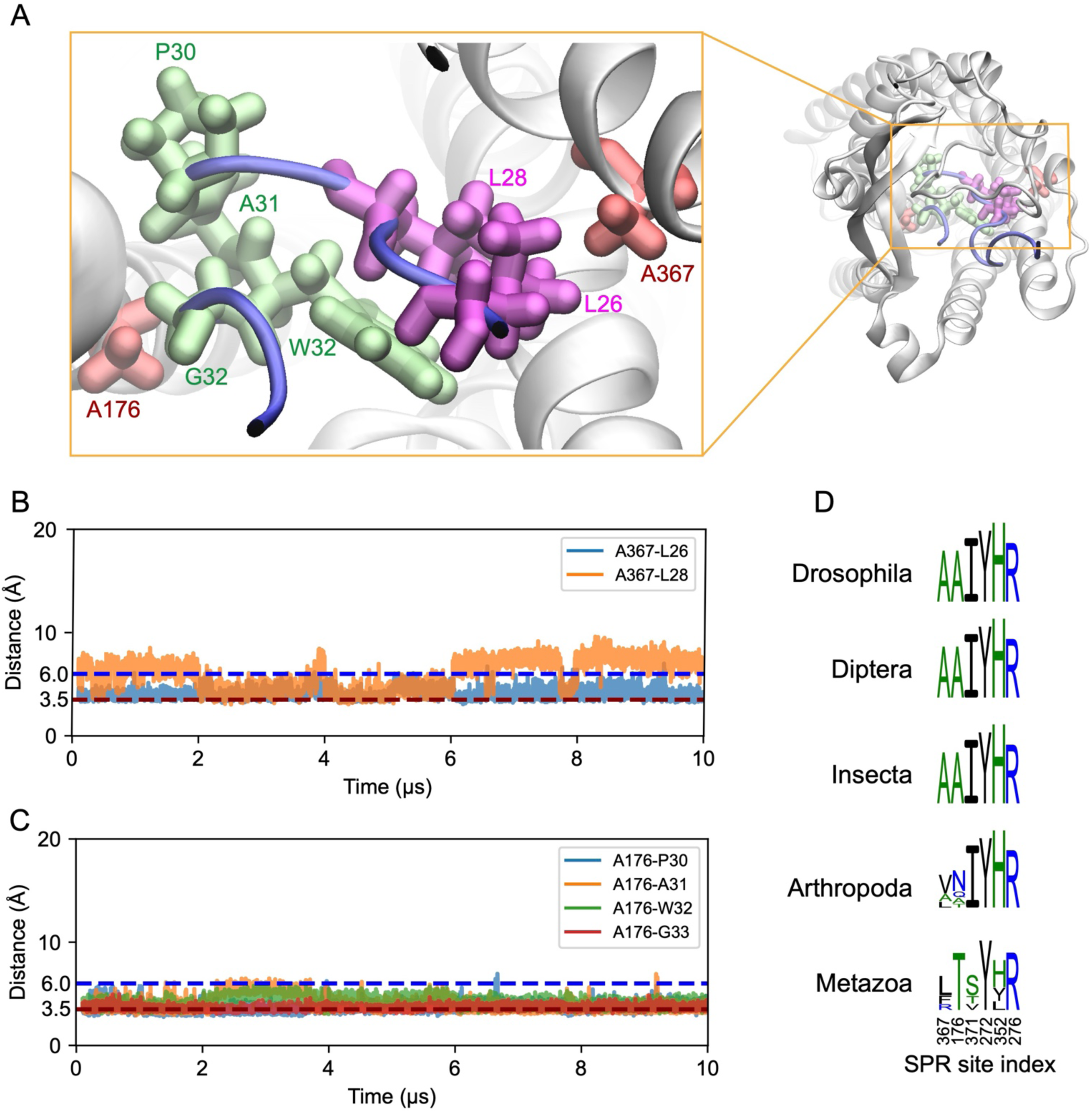
Pre-*Drosophila* mutations to A176 and A367 in D. melanogaster SPR (DmSPR) might create favorable conditions for the origination of SP-SPR interactions. (A) Structural overview of A176 and A367 in DmSPR and the frequently contacting residues within SP during MD simulations, including L26, L28, G29, P30, A31, W32, and G33. The structure is the representative conformation as shown in Figure 1E. (B) A367 interacts with L26 and L28 through hydrophobic interactions during MD simulations (Material and Methods). (C) A176 interacts with P30, A31, W32, and G33 through hydrophobic interactions during MD simulations (Material and Methods). (D) SPR sites 367, 176, 371, 352, and 276 in different ancestral sequences. Sites 367 and 176 showed the highest co-varying signals with SP binding states (Figure 4E). The two sites were highly conserved in *Drosophila*, *Diptera*, and *Insecta*. Sites 371, 272, 352, and 276 co-varying signals with SP binding states. The four sites were conserved across different clades.

## Discussion

### The evolutionary history of SP-SPR interaction

The sex peptide is one of the most well-studied seminal fluid proteins in nature. However, the sequence and functional conservation of the gene, along with its interaction with the SPR, have not been fully understood to date. In this work, we used a comprehensive set of computational tools to examine the interaction between the SP and SPR, as well as their origins. Our study revealed that SP has the potential to bind to the ancestral states of *Diptera* or even *Insecta* SPR (Figure 4). Indeed, an early study showed that SP could be a potent activator of SPR in moth (Yapici et al., 2008). This finding suggests that there were changes in SPR which could have allowed it to bind SP prior to SP’s speculated origin. This leads us to propose that these changes in SPR may have facilitated the origination and functional innovation of SP. This result is in line with previous findings that SP of *D. melanogaster* can have the ability to stimulate juvenile hormone synthesis and depress sex pheromone production in the moth *Helicoverpa armiger*a, which does not have a SP identified (Fan et al., 1999). Interestingly, besides SPR, several studies suggested it might be quite common for membrane receptors to appear early in evolution compared to their ligands across different species and taxa (Bridgham et al., 2006; Grandchamp & Monget, 2020; Ramilowski et al., 2015; Thornton, 2001).

### Changes in SPR predate the origin of SP

Additionally, we discovered that just a few amino acid modifications in SPR could potentially forge SP-SPR interactions. This discovery implies that the shift from a single MIPs-SPR interaction to a dual MIPs-SPR and SP-SPR interaction could have been a relatively uncomplicated evolutionary occurrence. Furthermore, we found that SP-SPR interactions predominantly occur through residues located at the MIP-SPR interface, which is a more ancestral function of SPR. This observation suggests that the co-evolution of the existing MIP-SPR interface may have presented opportunities for the emergence of novel interactions with new, or additional ligands, such as SP, when it appeared. Although AlphFold2 multimer prediction is generally highly accurate (Burke et al., 2023), our method currently has room for enrichment through the incorporation of biochemical and biophysical experiments – an avenue that presents an exciting prospect for future work. Recent studies have emphasized that AlphaFold2 has the potential to predict single mutation effects for protein monomers (Buel & Walters, 2022; McBride et al., 2023; Pak et al., 2023). In the context of this study, we posit that AlphaFold2 multimer prediction has the capability to predict how mutations in SPR might influence SP-SPR binding. To prove this, we carried out a *in silico* mutagenesis assay by generating random mutations in *D. melanogaster* SPR or SP sequence and performing AlphaFold2 multimer predictions using *D. melanogaster* SP and the mutated SPR sequences as well as using *D. melanogaster* SPR and mutated SP sequences (Material and Methods). We found that if random mutations occurred at the predicted SP-binding residues of SPR or C-terminal core regions of SP, the predicted SP-binding potential of the mutated sequences decreased drastically compared to the wild type *D. melanogaster* SPR (Figure S10). This highlights the potential and robustness of AlphaFold2 in our study. Overall, our findings shed light on a potential mechanism through which minor changes in the MIP-SPR interface could have created opportunities for a novel SP to interact and function. Note that the binding sites and interaction residues derived from our study are based on AlphaFold2 multimer predictions and MD simulations. A recent study showed that AlphaFold2 multimer predictions may not fully capture the influence of other variables on protein conformation and binding, but rather an exceptionally powerful hypothesis generator (Terwilliger et al., 2024), which is in line with the purpose of the current study. We anticipate future biochemical and biophysical studies to fully uncover the fascinating molecular details.

### SP and MIP share some interface with SPR – synergistic or antagonistic?

Our results show that SP and MIP share a common interface with SPR, while having some unique SP-SPR interactions. These interactions suggest a possible synergistic or antagonistic relationship between SP and MIP in the SPR interface, which is yet to be studied. However, it is important to note that SP and MIPS are expressed in different tissues and biological processes (Tsuda & Aigaki, 2016), and that MIP-SPR interactions originated earlier than SP-SPR interactions. This historical context suggests that the SPR initially evolved to accommodate MIP-SPR associations, which were later built upon by the emergence of SP-SPR interactions (Hopkins et al., 2024; Kim et al., 2010; Poels et al., 2010). The earlier establishment of MIP-SPR interactions might have set the foundation for the more complex network of interactions that we observe today.

When SP-SPR interactions occur, the immediate question that arises pertains to the fate of the MIP-SPR relationship. Does the SP-SPR interaction disrupt the MIP-SPR one, causing a conflict? Or is the issue effortlessly resolved because MIP generally exhibits tissue-specific expression different from SP? Given the tissue-specific expression of MIP, it’s plausible to suggest that the arrival of SP-SPR interactions might not drastically disrupt the existing MIP-SPR dynamic. Instead, it might lead to a more intricate network of interactions, each specialized for distinct tissue-specific functions.

Finally, the divergence of MIP sequences is particularly intriguing when examining the transition from *Arthropoda* to *Insecta* (Figure S11). By analyzing the differences in MIP sequence motifs (Nettling et al., 2015), we found that MIP sequences diverged much more significantly during the transition from *Arthropoda* to *Insecta* than from *Insecta* to *Drosophila* (Figure S11), indicating that the protein’s functions and interactions may have evolved in parallel. This divergence hints at adaptive changes to optimize the MIP-SPR interaction, reflecting the organism’s shifting needs and constraints over evolutionary timescales.

### The origin of SP may date back to the common ancestry of *Ephydroidea* or earlier

Previous results suggest that the sex peptide is a lineage-specific gene found in a subset of *Drosophila* species. However, our findings indicate that the sex peptide is more ancient than previously believed. The discovery of this gene in *S. lebanonensis* suggests that it may be conserved across all *Drosophila* species, despite occasional gene loss in some. This is in line with the recent findings from Hopkins et al. (Hopkins et al., 2024). The presence of the gene in other flies, such as the melon flies *Zeugodacus cucurbitae*, remains uncertain as the potential candidate gene in *Z. cucurbitae* lacks a signal peptide (Figure S12). In vinegar fly *Phortica variegata*, we identified a sequence that, while lacking a signal peptide, contains two W(X)_11_W motifs in the middle (Figure S13). Since the W(X)_11_W structure is not at the C-terminal, and the sequence length between the two Ws is 11 amino acids, rather than the typical 8, we argue that this might not be a sex peptide protein. It could represent the ancestral form of the sex peptide, or an unrelated gene with a similar W(X)_n_W motif. In either case, it appears that the sex peptide may have originated around the time of divergence between these species. So far, there is no solid evidence to suggest the sex peptide exists in mosquitoes or moths (Tsuda & Aigaki, 2016). However, we were able to discover a bunch of possible W(X)_n_W motifs in their genomes. In our study, we only assigned a SP gene if it followed a strict SP gene structure pattern, i.e., the two exons and one intron pattern observed in *Drosophila* species. A limitation would be that if the gene in older lineages evolved to have new gene structures, we could fail to identify orthologs in older lineages.

### The origin of SP - *de novo* or divergence?

The SP is produced in male accessory glands of *D. melanogaster* and plays a crucial role in the post-mating behavioral and physiological responses in female fruit flies. Despite its importance in some fruit fly species, the gene is not found in other lineages (Tsuda & Aigaki, 2016). This observation sparks a fascinating discussion about its origin and evolutionary path. In principle, the pattern could be explained by two main theories: *de novo* origin and divergence.

*De novo* gene birth refers to the process where new genes arise from non-genic DNA sequences (Begun et al., 2006, 2007; Levine et al., 2006). These genes do not have any ancestral gene copies, which can explain their absence in other species. Instead, they evolve from previously non-genic sequences that gain function, usually due to changes in the regulatory regions that start to drive the expression of these sequences in a context where they can impact fitness. This phenomenon has been recognized in various species and is considered an important source of evolutionary novelty. Although hard to prove, SP could indeed be a product of *de novo* gene birth. If true, it suggests that at some point in the evolution of the fruit fly lineage, a non-genic sequence gained function, leading to the formation of SP.

Two pieces of evidence support this idea. First, there is evidence that accessory gland proteins may be easily originated *de novo* (Peng & Zhao, 2024). In fact, the first several works identifying *de novo* genes by the Begun group (Begun et al., 2006, 2007; Levine et al., 2006) were primarily accessory gland proteins. Additionally, several other *de novo* genes have also been shown to be involved in *Drosophila* fertility (Gubala et al., 2017; Lange et al., 2021; Reinhardt et al., 2013; Rivard et al., 2021). Second, although SP has a critical function in *D. melanogaster*, various studies suggest that it is less important in some other *Drosophila* species (Tsuda et al., 2015). This is in line with the simple assumption for novel gene origination, that a new gene is born with a function that can be selected by natural selection. Over time, the new gene may be recruited to complex functional networks and generate pleiotropy.

How likely is it that a gene similar to sex peptide could be born *de novo*? The sex peptide contains a signal peptide, a middle part that functions in sperm binding and juvenile hormone response, and a conserved C-terminal region that binds to SPR. Signal peptides, similar to transmembrane sequences, can be easily formed by hydrophobic sequences (Aspden et al., 2014; Tassios et al., 2023). The conserved C-terminal function W(X)_8_W partly relies on the two tryptophan residues (Kim et al., 2010). We asked, how likely intergenic sequences would produce a sequence with two tryptophan residues and several other amino acids in between. The answer is that there are thousands of such sequences in intergenic sequences (Figure S14). Thus, when opportunities arise, if these sequences are transcribed or translated in the right cell types, they may produce a function that interacts with the preexisting SPR.

Alternatively, the lineage-specific pattern could be a result of divergence. Divergence here refers to the process in which an existing gene evolves beyond recognition, i.e., it changes so much over time that it is impossible to identify it as the same gene in different species. Given that the SP is relatively short (36 amino acids), and assuming little to no selective pressure maintaining its sequence, it could have diverged significantly from its ancestral state, rendering it unrecognizable in other species. In addition to genetic drift caused by near neutral processes, natural selection and positive selection can also cause such patterns. If this is the case, it suggests a scenario where the SP or its predecessors existed in other lineages (e.g., mosquitoes, moths, butterflies), but over time, due to lack of constraint, it has changed to the point of being unique to fruit flies.

How likely is a protein to become unrecognizable after a certain period of time? For a long protein, the probability is low. However, for a short protein, it is entirely possible. Overall, signal peptides have a lower sequence constraint, and their substitution rate is three to four times higher than that of mature proteins (Canale et al., 2018). Given a timeframe longer than 60 million years, it is feasible for signal peptides to change without becoming recognizable (Venkataraman et al., 2023). As for C-terminal sequences, speculation becomes more challenging. Even for a gene with entirely different sequences, they still need to bind to SPR. SPR has a pocket that can only bind to certain sequence motifs, making it hard to envision a scenario where sequences change completely but can still bind to SPR. Future structural modeling might offer new insights into the binding potential of SPR. Currently, tools like AlphaFold2 can be used to predict interaction affinity between SPR and a random amino acid. However, executing the probability would require a scale of 20^10, which is not feasible in a short time frame but might be straightforward in the future. Alternatively, there might soon be a way to predict all the possible binding sequences of a protein pocket - such a method does not exist yet, but recent computational methods such as large language models (Lin et al., 2023) or building ultra large candidate libraries (Lyu et al., 2019), could make such an idea possible. This advancement could help us answer whether genes with completely different sequences can still serve the same role in functional networks.

In conclusion, both the *de novo* origin and divergence theories provide plausible explanations for the apparent lineage-specificity of the SP gene. It is also possible that both mechanisms have played a role to some extent. Determining which one is the most probable would necessitate a more profound examination of the evolutionary history of the SP gene. This could be achieved through advanced computational analyses and phylogenetic studies, aided by the generation of high-quality genomes and annotations in a variety of non-*Drosophila* insect species.

If SP turns out to be a lineage-specific new gene, then studying its origin and interaction with SPR also provides an intriguing example of how a novel ligand can quickly incorporate into a functional network and become functionally important. Given that there are many novel and lineage-specific ligands in various species (Ramilowski et al., 2015), including humans, understanding the origin of SP and the SP-SPR interaction provides fundamental insight into ligand-receptor evolution and the evolution of complex intercellular communications.

## Material and Methods

### AlphaFold2 mulitmer predictions

We used AlphaFold2 multimer (Evans et al., 2022), version 2.2.2, with default settings for all the multimer predictions in this study. For each prediction, we extracted interface pTM (ipTM) and predicted aligned error (PAE) from the first ranked model. We adapted the scripts from Burke et al (Burke et al., 2023) to compute pDockQ value using a cutoff of 7 Å. To gain a better understanding of the predictions, we plotted in Figure S1D-S1E the statistics of the multiple sequence alignments and PLDDT curves from the AlphaFold2 multimer prediction of *D. melanogaster* SP-SPR.

### Identification of potential SP genes

We used the partial sequence of *D. melanogaster* SP as the query and performed a *tblastn* (Altschul et al., 1990) search against NCBI eukaryotes representative genomes (Sayers et al., 2022), which contains high quality genome assemblies of 1642 eukaryotic species. Similar to a recent study by Hopkins et al (Hopkins et al., 2024), we used the most C-terminal sequence of SP, KWCRLNLGPAWGGRC, since the C-terminus is highly conserved, while N-terminus is highly diverged (Figure 1). We collected possible similar sequences with an arbitrarily large *tblastn* E-value cutoff of 9999 in order to include as many *tblastn* hits as possible. At this step, there are 191 *tblastn* hits with query coverage ranging from 60% to 100% and sequence identity ranging from 47% to 100%. We then used a gene structure aware spliced aligner, SPALN (Iwata & Gotoh, 2012), to verify if the hits could potentially be SP genes. We assigned a *tblastn* hit to a potential SP gene only when SPALN predicted a regular SP gene structure with canonical start/stop codons and no frameshifts. A regular SP gene in *Drosophila* species means that it contains two exons and one relatively short intron. In the *Drosophila* species, the length of the short intron ranges from ∼30 to ∼70 bp. While for the two exons, the length was ∼80 to ∼130 bp for the first exon, and ∼50 to ∼60 bp for the second exon. Following this step, we were able to identify 61 potential SP genes across 33 species. We used MAFFT (Katoh et al., 2002) to align the DNA sequences and protein sequences. We further visualized the alignments by MVIEW (Brown et al., 1998) and sequence motifs by WebLogo (Crooks et al., 2004). We then used iqtree2 (Minh et al., 2020) to infer the phylogenetic relations of the 61 SP genes. We further performed SignalP6.0 (Teufel et al., 2022) analysis to predict whether the SP proteins contain N-terminal cleavage signals.

### Accessory gland mass-spectrometry

Mixed-age accessory glands from both *D. suzukii* and *D. subpulchrella* were dissected and placed on ice-cold 1X PBS. Each set of accessory glands was then transferred to a volume ranging from 50-110 μl of lysis buffer, homogenized, and the resulting mixture was collected. Proteins were precipitated using ice-cold acetone. Acetone was removed and the pellets were dissolved in 8M Urea/50mM/20mM DTT ammonium bicarbonate and incubated at room temperature, alkylated (iodoacetamide), and digested with Endoproteinase LysC at <4M urea for 4 hours. Hereafter, the samples were diluted to <2M urea and trypsinized overnight. Peptides were micro-solid phase extracted (Rappsilber et al., 2003) and analyzed by LC-MS/MS using either a 90-minute gradient separated using a 12cm 100um/15cm packed emitter or a 120-minute gradient separated using an EasySprayer (75um/25cm) column. Data were recorded in high resolution/high mass accuracy mode using either a QExactive-HF or a Fusion Lumos. For each sample set, a 2nd run was designed to target several copies of sex peptides using Parallel Reaction Monitoring (PRM) (Peterson et al., 2012). The data were processed using ProteomeDiscoverer v1.4 or v2.5 and searched (Mascot) against a custom database concatenated with LysC and trypsin. Tandem MS spectra of selected matched peptides were compared to in-silico calculated fragmentation (Prosit (Original model, 2019) (Gessulat et al., 2019).

### Molecular dynamics simulations

We used AlphaFold2 to predict the structural model of the *Drosophila melanogaster* SP-SPR complex and MIP-SPR complexes as described in section *AlphaFold2 mulitmer predictions* in *Methods*. We used CHARMM-GUI membrane builder (Jo et al., 2008; J. Lee et al., 2016; Wu et al., 2014) to build a heterogeneous bilayer membrane environment for the complexes with cholesterol (30%), POPC (35%), and POPE (35%) lipids (Figure S4). The system was then set up by the additive CHARMM36 protein force field. We added TIP3P water with a length of 12 Å to the top and bottom of the system. We then added additional Na^+^ and Cl^-^ ions to neutralize the system and to reach an ion concentration of 0.15 mM. We further minimized the potential energy of the system by the steepest descent method followed by the conjugate gradient method until the maximum force was smaller than 200 kJ mol^-1^ nm^-1^. We then used GROMACS (Abraham et al., 2015) to perform subsequent MD simulations. For the SP-SPR complex, we carried out a 10-μs and two additional 2-μs MD simulations. For each of the five MIP-SPR complexes, we carried out a 4-μs and two additional 2-μs MD simulations. The simulation trajectories were recorded every 100 ps. For each frame of each trajectory, we used getcontacts (getcontacts.github.io) to extract interactions between SP or MIP peptides and SPR with the option “--itypes all --vdw_epsilon 1.5”. We then computed the frequency of each interaction during the MD simulations. For each of the SP-SPR and MIP-SPR complexes, we considered two residues to be interacting only when the frequency of their interaction is greater than 5% and frequently interacting only when the frequency is greater than 50% in all the three replicate MD simulations.

### Ancestral sequence reconstruction

We searched through orthoDB (Kuznetsov et al., 2023) to obtain SPR genes and orthologs. We first collected 328 SPR protein sequences in 228 *Metazoa* species. We used MAFFT (Katoh et al., 2002) to align the sequences. We further excluded the sequences or fragments that have less than 50% coverage compared to SPR gene in *D. melanogaster*. Finally, we obtained 254 SPR genes across different *Metazoa* species. We then used iqtree2 (Minh et al., 2020) to construct the SPR gene tree. We used the maximum likelihood method in PAML (Yang, 2007) to infer the ancestral sequences of SPRs in different clades, including *Drosophila*, *Diptera*, *Insecta*, *Arthropoda*, and *Metazoa* species. We extracted the most probable ancestral sequences, and alternative states with probability greater than 5%.

### In-silico SP-SPR interacting pattern co-varying analysis

For each of the *Drosophila*, *Diptera*, *Insecta*, *Arthropoda*, and *Metazoa* clades, we generated 100 alternative ancestral sequences of SPR. For each of these alternative ancestral SPR sequences, we randomly selected half of the sites with alternative states and mutated the selected sites to one of their alternative states. By doing this, we aimed to avoid almost identical alternative ancestral SPR sequences, which could be challenging for AlphaFold2 (Buel & Walters, 2022). We then used AlphaFold2 (Evans et al., 2022) to predict whether the alternative ancestral SPR sequences could bind *D. melanogaster* SP. We used the pDockQ metric to determine whether SP could bind or not bind to ancestral SPRs. The pDockQ metric has been applied to distinguish interacting from non-interacting proteins (Bryant et al., 2022) and to structurally resolve human PPI network (Burke et al., 2023). Here, we defined SP binding state of an ancestral SPR sequence to be true if the corresponding AlphaFold2 prediction has a pDockQ value greater than 0.3, which is more stringent than the proposed cutoff value of 0.23 (Bryant et al., 2022). Whereas if the pDockQ value was smaller than 0.23, we defined its SP binding state to be false. We extracted the SPR sites that had contacts with either SP or MIP peptides in the MD simulations with frequencies greater than 5%. We took the advantage of Gremlin, a program that could accurately capture co-evolution patterns in protein families (Ovchinnikov et al., 2014), to infer the residues that co-varied with the SP binding state of ancestral SPRs. A simplified scheme of the analysis can be found in Figure 4D. Specifically, we constructed a pseudo alignment, [X_1_, X_2_, …, X_n_; X_n+1_], from the extracted SPR sites, where X_1_ to X_n_ (n=64) represent the predicted SPR interface as described above, while X_n+1_ represents the predicted binding potential of the corresponding alternative ancestral sequence. We arbitrarily used “H” to stand for potentially binding events and “P” to stand for potentially non-binding events. We then used Gremlin (Ovchinnikov et al., 2014) to compute the co-varying scores between X_n+1_ and other sites from the predicted SPR interface using a maximum likelihood method.

### AlphaFold2 predictions of SP-SPR interactions with in silico random mutations in SP or SPR

To investigate whether AlphaFold2 could predict how mutations in SPR affect SP-SPR interactions, we generated 100 mutated SPR sequences, each with 30 random mutations. We chose 30 random mutations to mimic the minimum number of sites that have alternative ancestral states in each of the *Drosophila*, *Diptera*, *Insecta*, *Arthropoda*, and *Metazoa* clades. We used AlphaFold2 to predict the interactions between SP and the mutated SPR sequences. We also checked whether the mutations were in the SP binding interface. Meanwhile, to investigate whether AlphaFold2 could predict how mutations in SP affect SP-SPR interactions, we generated 200 mutated SP sequences. Specifically, the 200 mutated SP sequences include 50 with 3 random mutations in the N-terminal regions, 50 with 6 random mutations in the N-terminal regions, 50 with 3 random mutations in the C-terminal core regions, and 50 with 6 random mutations in the C-terminal core regions. We used AlphaFold2 to predict the interactions between SPR and the mutated SP sequences.

### Identification of the W(X)nW motif in the *D. melanogaster* genome

We used ORFfinder (Sayers et al., 2022) to search for all the intergenic ORFs in the *D. melanogaster* genome by a six-frame translation of all the intergenic regions. We then translated the ORFs and searched for W(X)_n_W motif in the translated sequences. We counted the length of the motif and the number of motifs in each of the ORFs. We were able to identify thousands of W(X)nW motifs with the motif length n of 8 to 9. Some of the sequences have high sequence identity compared to the W(X)nW motif in MIP peptides and SP (Figure S14).

## Data availability

Primary data have been uploaded to PRIDE database (www.ebi.ac.uk/pride/) with with project accession: [PXD044633], username, and password: LKWAT1Qk. The top ranked predicted structural models of SP-SPR complex and five MIP-SPR complexes along with the MD simulation trajectories are available at Figshare (https://doi.org/10.6084/m9.figshare.25245223.v1). Code and scripts for this study are available on GitHub (https://github.com/LiZhaoLab/DrosophilaSexPeptide).

## Author contribution

J.P. and L.Z. conceived the study, designed the experiments, and formulated the analyses. J.P. conducted all analyses except those noted below. N.S. identified gene duplications in the suzukii species complex and conceived mass-spectrometry analysis. H.M. analyzed the mass-spectrometry results. J.P. and L.Z. authored the paper.

## Acknowledgement

We thank Christopher B. Langer and Vivian Yan for dissecting the accessory glands. We thank Jason Banfelder, Bala Jayaraman, and The Rockefeller University High Performance Computing (HPC) Center for their support in computation. We are grateful to Dr. Mariana Wolfner for critically reading the manuscript and providing valuable feedback. The authors also thank Drs. Ben Hopkins and Artyom Kopp for their comments and suggestions on a previous version of the manuscript.

