## Supplementary figures for "The Origin and Evolution of Sex Peptide and Sex Peptide Receptor Interactions"

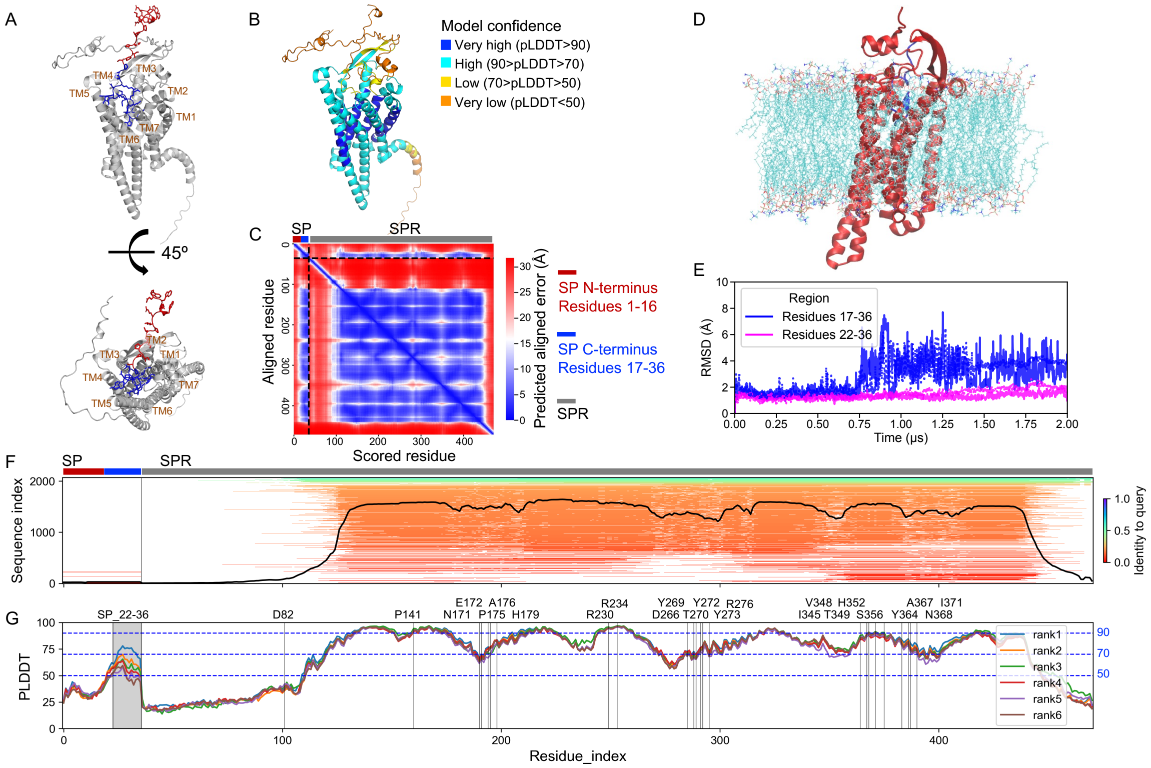

Figure S1. AlphaFold2 prediction of the SP-SPR complex. (A) Predicted complex structure. (B) Predicted complex structure colored by PLDDT. The corresponding PLDDT curve is shown in (G). (C) Predicted aligned error between SP and SPR in AlphaFold2 prediction, where blue regions indicate residues with high confidence and small expected positional error while red regions indicate residues with low confidence and large expected positional error. (D) Configuration of the bilayer membrane environment for the SP-SPR complex. (E) Time resolved RMSD curves from the two additional 2-μs MD simulations (one is shown in solid line and the other in dashed line) of the SP-SPR complex. Statistics of multiple sequence alignments (F) and PLDDT (G) obtained from AlphaFold2 multimer prediction of *D. melanogaster* SP-SPR interaction.

A

```
     M  Q  A  V  A  I  I  F  F  F  L  L  G  V  G  L  V  -  -  - | SP (Sleb)
  1 ATGCAAGCTGTCGCAATCATTTTTTTCTTCCTACTAGGAGTTGGTCTAGTC---------| sp (Sleb)
  1  M  K  T  L  A  L  F  L  V  L  V  C  V  L  G  L  V  Q  A  W | SP (Dmel)

     -  -  -  -  -  -  -  -  -  -  S  J  N  P  A  T  J  A  R    | SP (Sleb)
 51 ------------------------------TTGAGCAACCCCGCTACAAGTGCTAGAGgt| sp (Sleb)
 21  E  W  P  W  N  R  K  P  T  K  F  P  I  P  S  P  N  P  R    | SP (Dmel)

                                                      D  K  | SP (Sleb)
 81 aacgaaataccaactaattagaaaacgcttataaaaatataaaatttatttcagATAAAT| sp (Sleb)
 40                                                   D  K  | SP (Dmel)

    W  C  R  L  N  L  G  P  V  W  G  G  R  C                    | SP (Sleb)
141 GGTGCCGCCTGAATTTGGGCCCGGTCTGGGGTGGCAGGTGCtaa                | sp (Sleb)
 42 W  C  R  L  N  L  G  P  A  W  G  G  R  C                    | SP (Dmel)
```

B

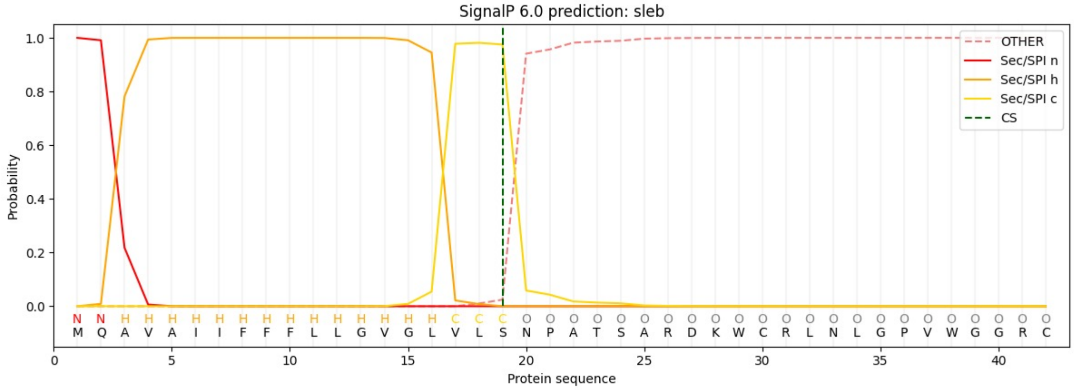


C

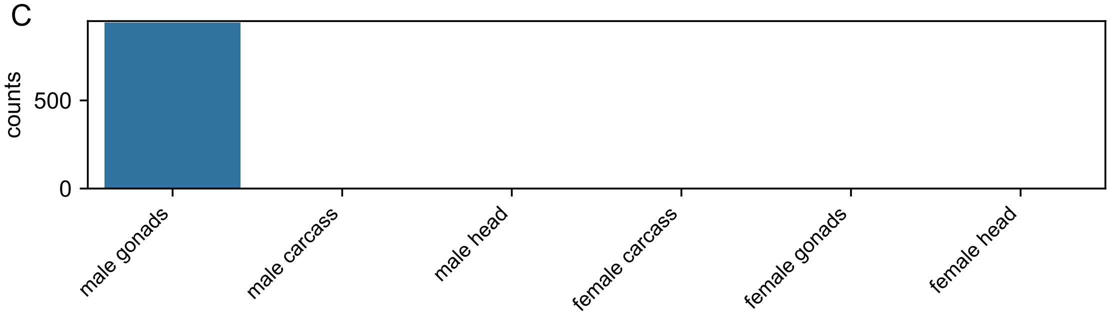


Figure S2. The *SP* gene in *S. lebanonensis*. (A) The gene structure as predicted by SPALN. (B) The SP of *S. lebanonensis* contains a signal peptide at the N-terminus. (C) The SP of *S. lebanonensis* shows male gonad-specific expression.

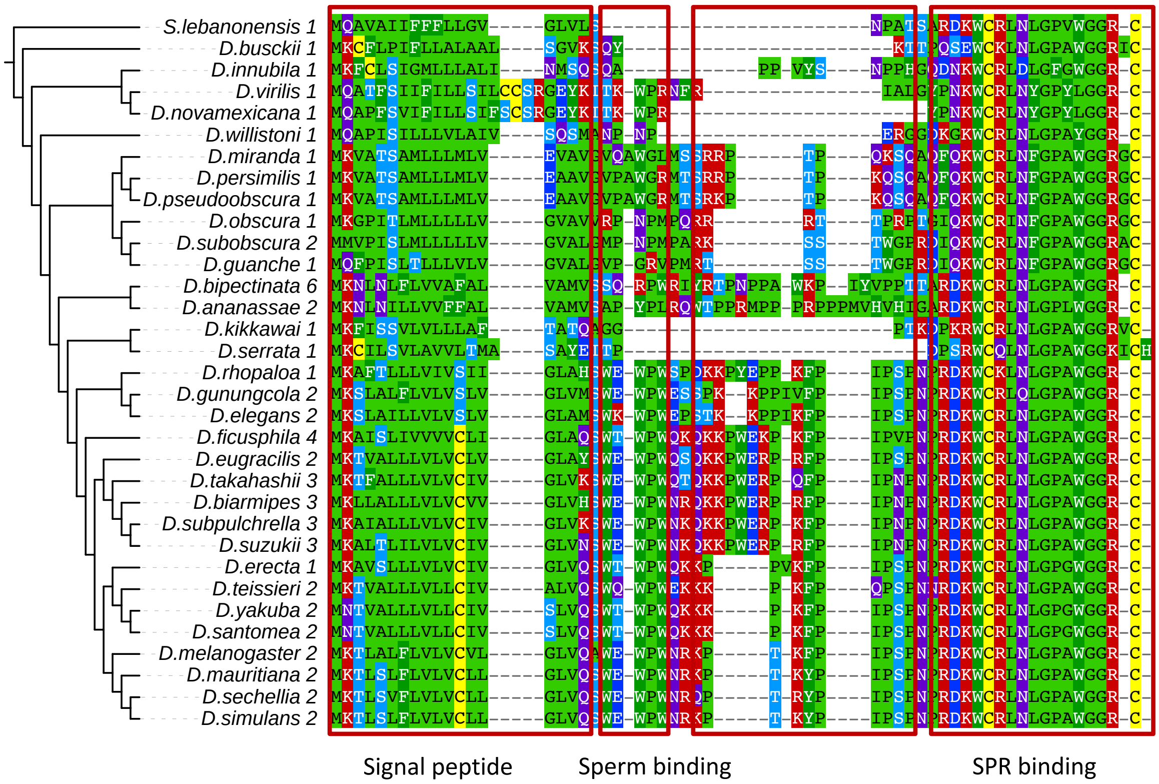


Figure S3. Multiple sequence alignments of representative SP proteins in the identified species.

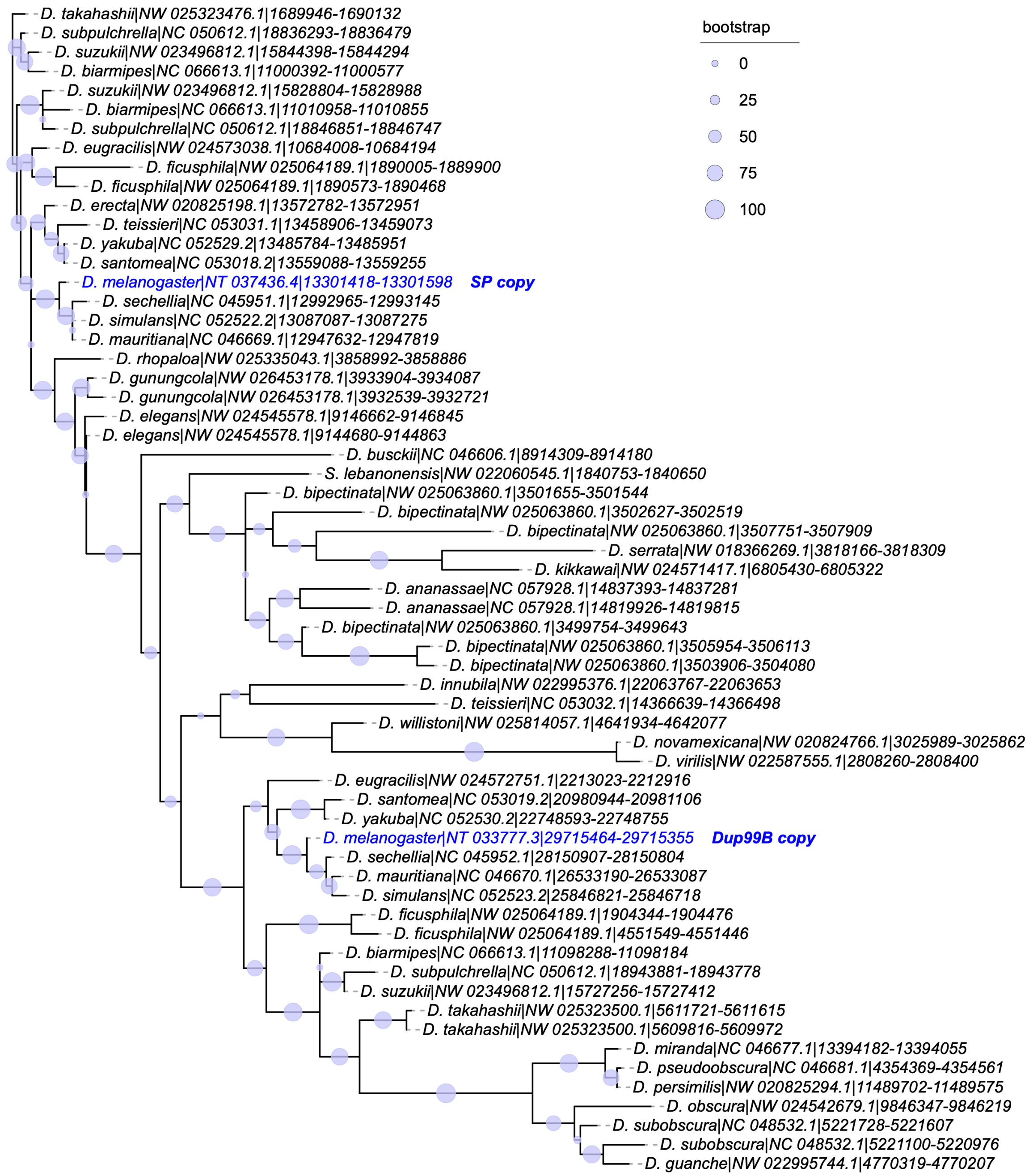


Figure S4. Phylogenetic tree of the 61 identified *SP* genes. The two copies in *D. melanogaster*, named as *SP* and *Dup99B*, are highlighted in blue.

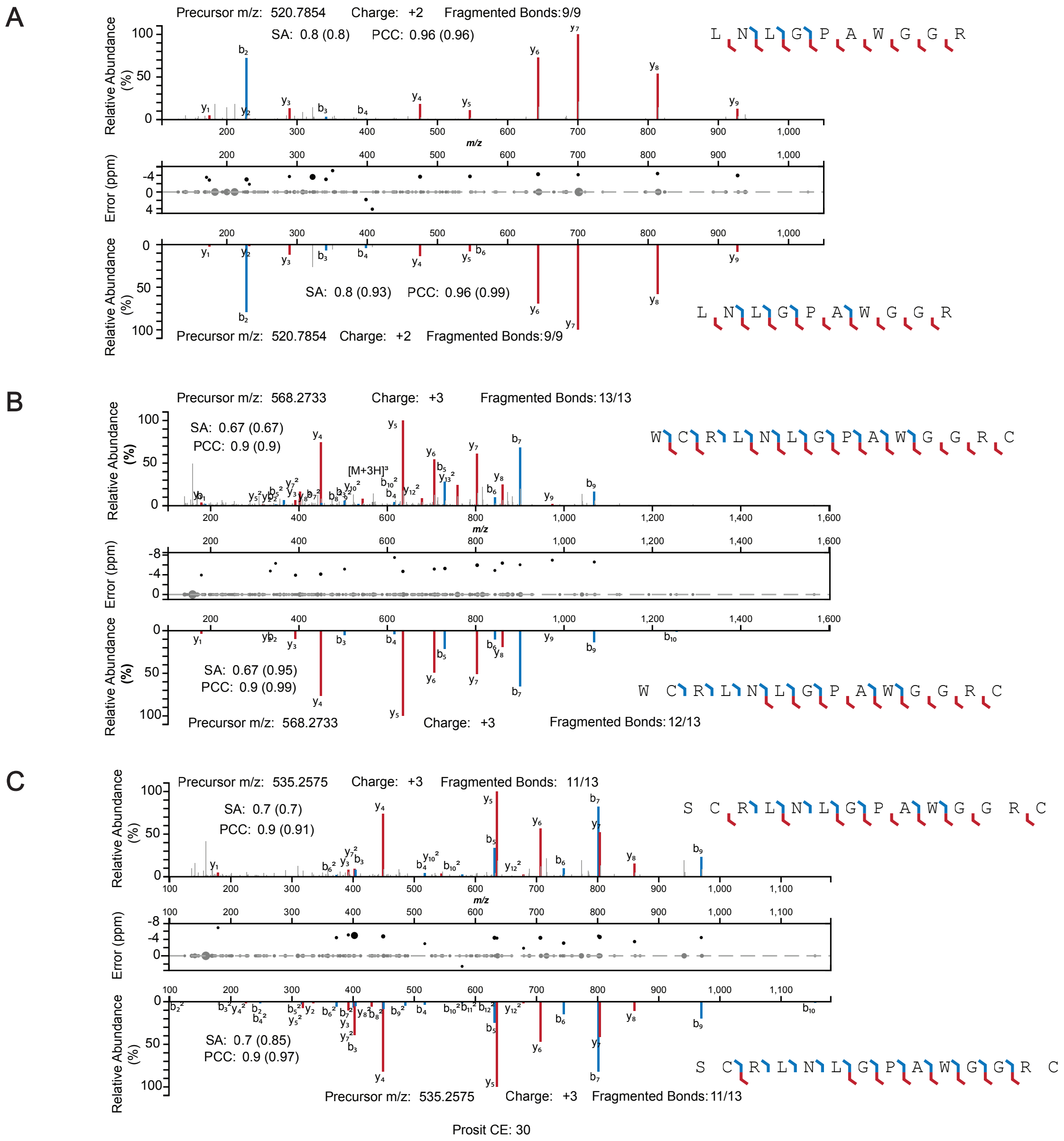


Figure S5. Example peptides supporting the translation of different copies of sex peptide. Acquired tandem MS high-resolution spectra (30,000@200 Th) (upper panel of A, B, and C) are matched to the peptides in *D. subpulchrella* and *D. suzukii*, respectively. A: LNLGPAWGGR[2+], Mascot score: 60, shared by XP_037720360.1, LOC119553811, and XP_037721024.1 in *D. subpulchrella*, B: WCRLNLGPAWGGRC[3+], Mascot score: 30, specific to XP_016924329.1 in *D. suzukii*, and C: SCRLNLGPAWGGRC[3+], Mascot score: 28, specific to XP_016924330.1 in *D. suzukii*. The observed spectra (in the upper panel of A, B, and C) are compared with in-silico fragmentation calculated by Prosit (Original model, 2019), listed in the bottom panel of A, B, and C. Cysteines were treated as carbamidomethylated.

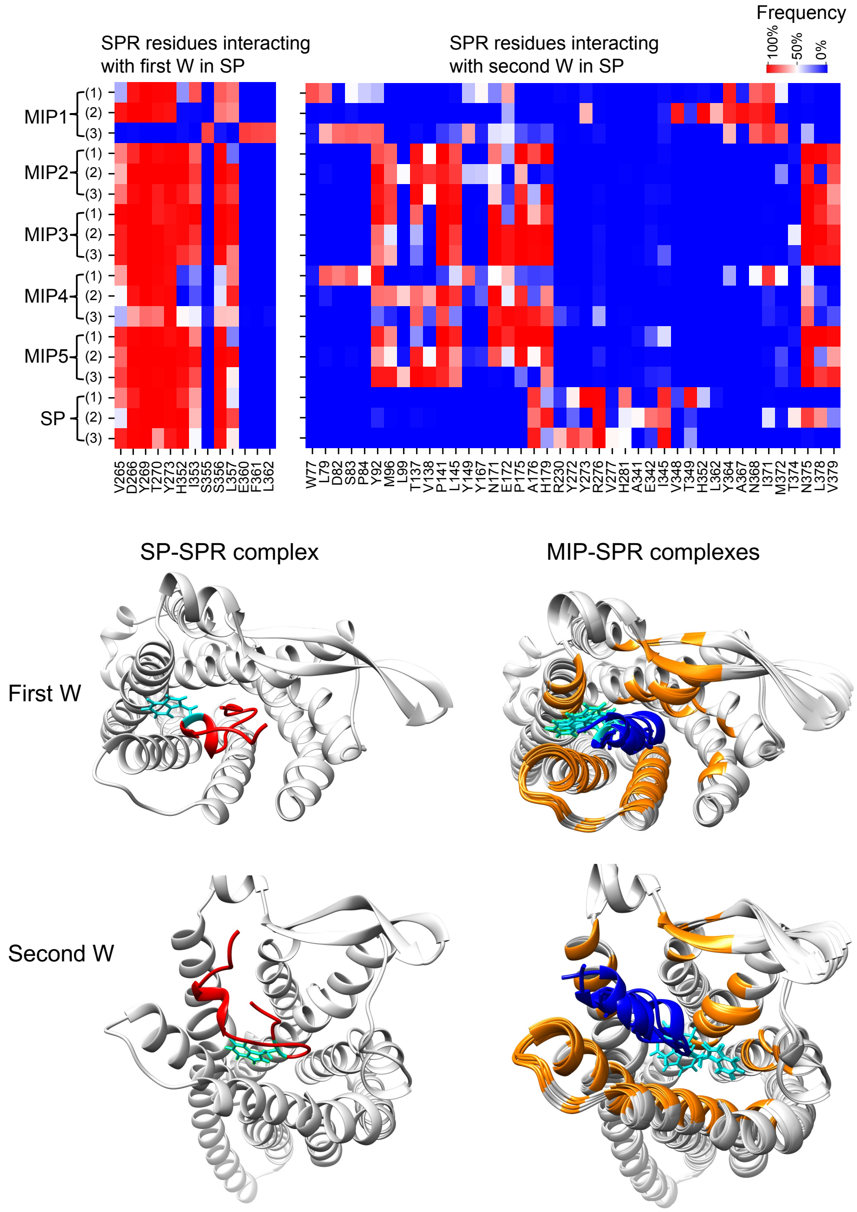


Figure S6. Predicted interaction patterns of the two conserved tryptophan residues of the W-(X)n-W motifs in SP and MIP peptides. Top panel: Heatmap of interaction frequencies of the two tryptophan residues in MD simulations. The first W residues shared similar interaction patterns across SP and MIP peptides, while the second W residues displayed differences. Middle panel: Detailed close-up view of the position of first W residues in SP-SPR complex (left) and MIP-SPR complexes (right). Bottom panel: Detailed close-up view of the position of second W residues in SP-SPR complex (left) and MIP-SPR complexes (right).

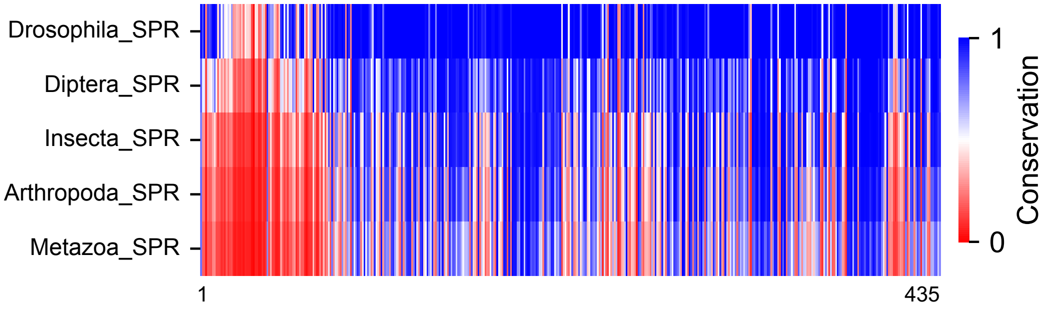


Figure S7. Conservation of SPR sequences across *Metazoa* species. The N-terminal of SPR is highly diverged while other regions can be conserved.

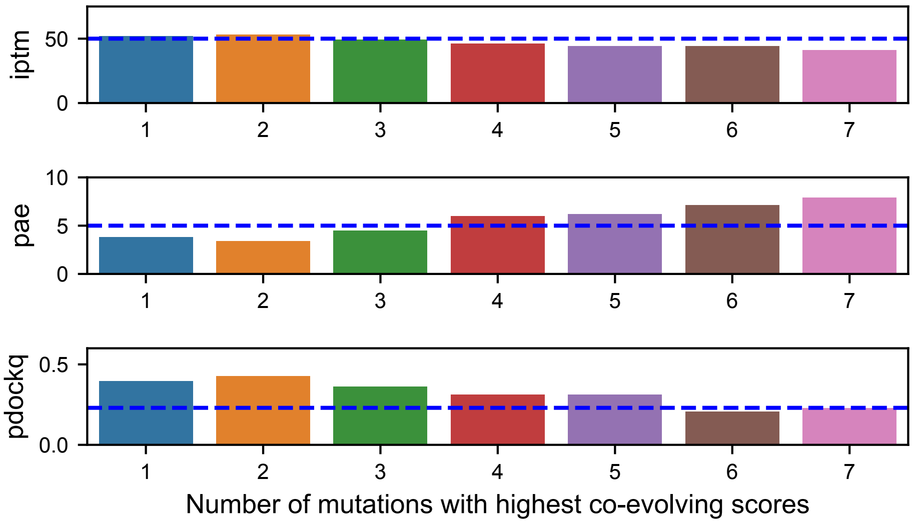


Figure S8. AlphaFold2 predictions of SP-SPR interactions by changing SPR different number of residues with top co-evolving scores with SP binding potentials back to their most ancestral states. The SP binding potential became not acceptable when top 6 scored residues were mutated (Material and Methods), indicating the top-scored residues alone could potentially switch on/off SP-SPR interactions.

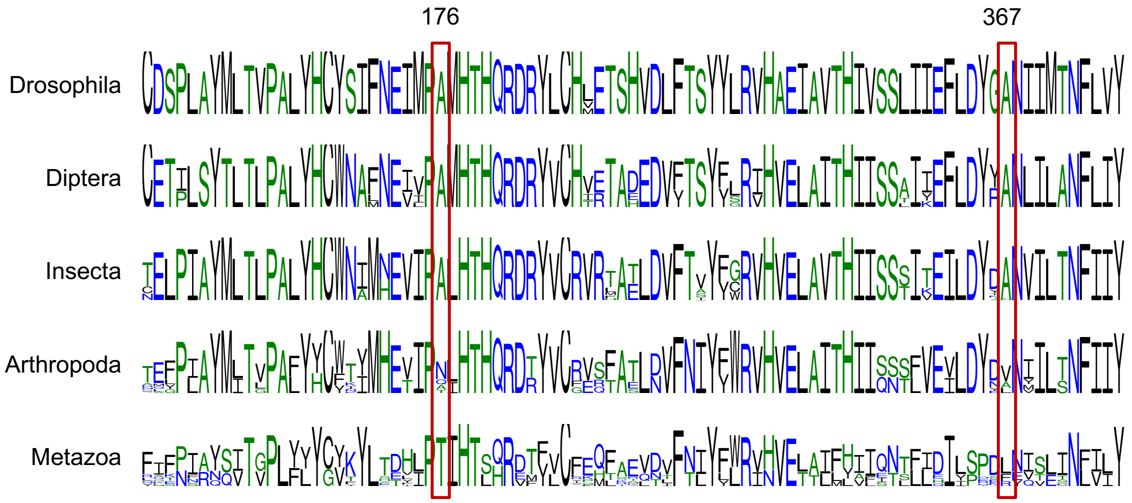


Figure S9. Sequence logo of ancestral SPR residues that interact with SP and MIP in different clades.

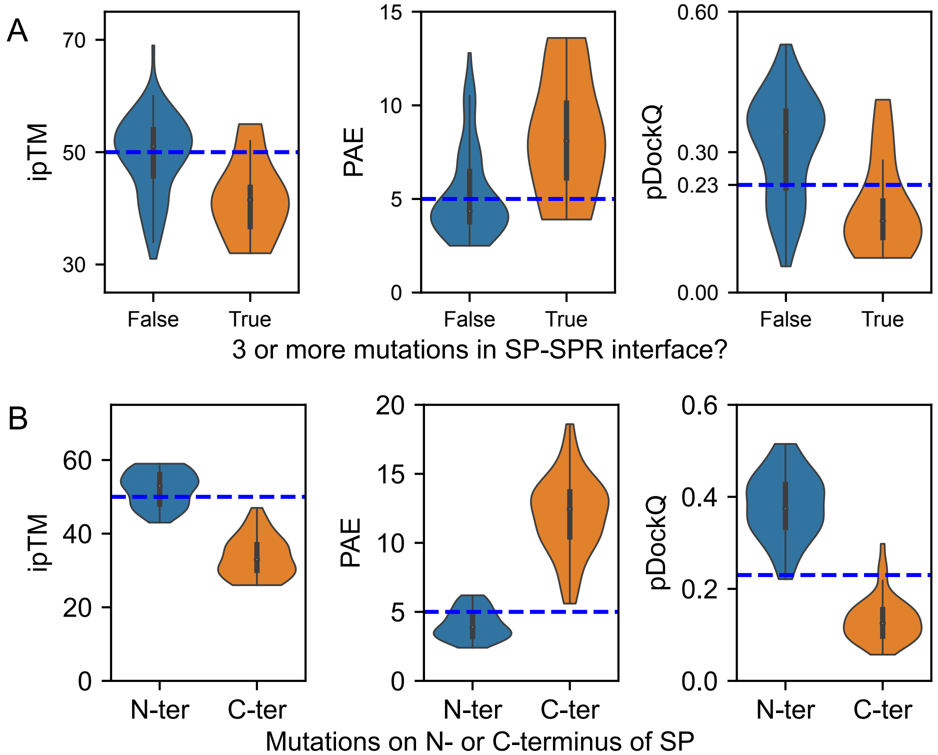


Figure S10. AlphaFold2 could predict whether mutations in SP and SPR affect SP-SPR binding. (A) The interface TM-score (left panel), PAE (bottom panel), and pDockQ (right panel) of random SPR mutations suggested SP-SPR interactions decrease significantly when the random mutations contain SP-SPR interface residues. (B) The interface TM-score (left panel), PAE (bottom panel), and pDockQ (right panel) of random SP mutations suggested SP-SPR interactions decrease significantly when the random mutations happen on the C-terminus of SP, which was predicted to be important for SP-SPR interactions in this study.

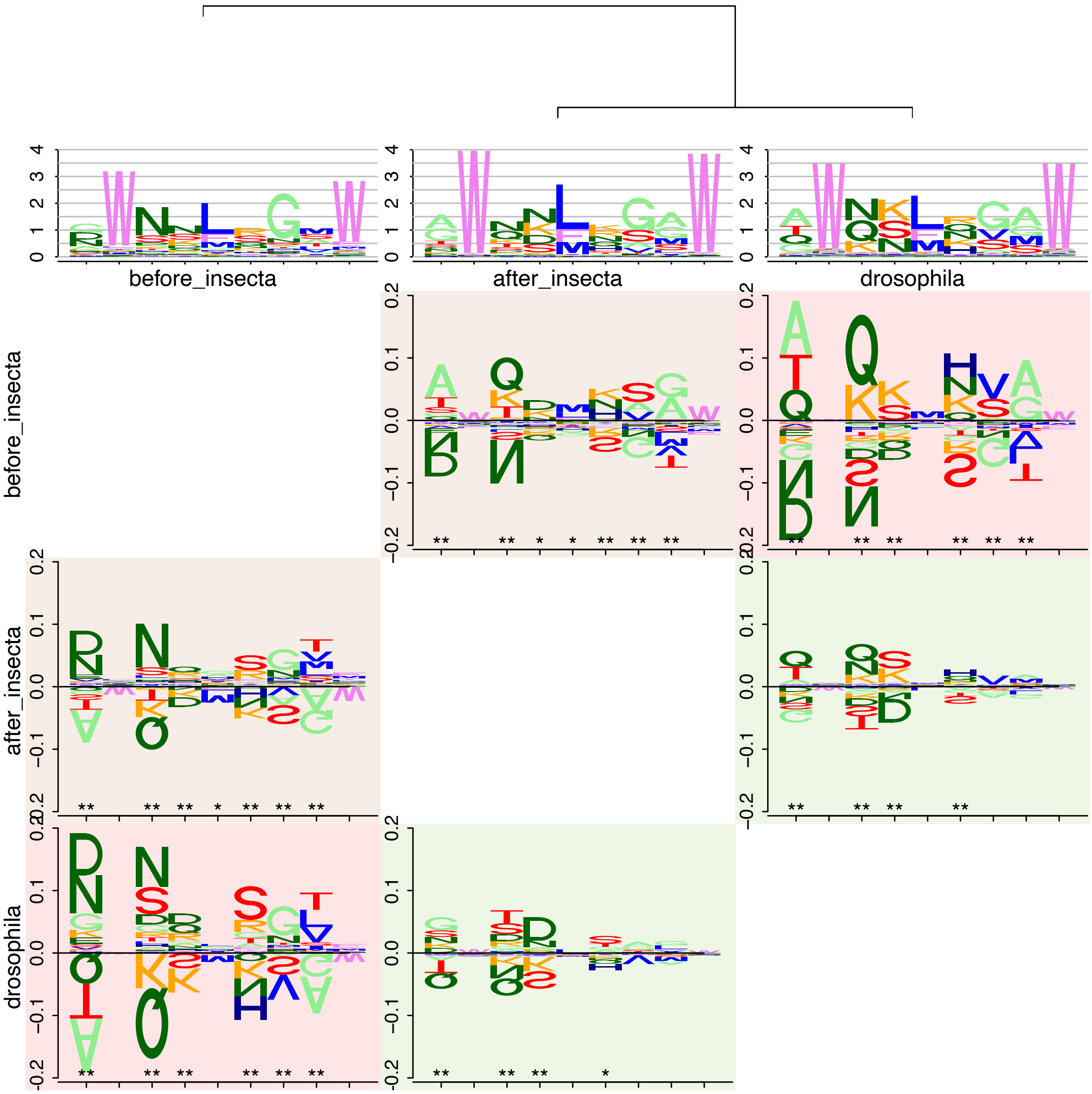


Figure S11. Sequence logo of MIP peptides in *Drosophila*, in taxa before *Insecta*, and in taxa after *Insecta*. The “before *Insecta*” MIP motif showed significant differences from the “after *Insecta*” and “in *Drosophila*” motifs, with almost all sites except for the two W amino acids being significantly different. Compared to the large differences between “before *Insecta*” and “after *Insecta*”, the differences between “after *Insecta*” and “in *Drosophila*” motifs were smaller.

```
  M  G  K  E  C  L  E  L  S  N  A  A  L  F  P  E  P  F  N  V
 ATGGGAAAGGAGTGCTTGGAACTTTCGAATGCGGCACTGTTTCCGGAACCTTTCAATGTG| zcuc
  -  -  -  -  -  -  -  M  K  T  L  A  L  F  L  V  L  V  C  V | SP

  K  E  Y  Y  E  P  L
 AAGGAATACTATGAACCATTGgtgaaaggtataatatcttacattatgcattatatataa| zcuc
  L  G  L  V  Q  A  W                                         | SP

;; skip 300 nt's
                                           K  W  -  W  Q  N  C
 gcattgtgcatgtttcaaaggcaacaactatagcacgagAAATGG---TGGCAAAATTGC| zcuc     Not a signal peptide
                                           E  W  P  W  N  R  K | SP

  R  J  L  Y  -  -  -  -  -  -  -  -  Q  D  R  W  C  K  L  P  Y
 AGGAGCTTATAC---------------------CAAGACAGATGGTGCAAATTGCCATAC| zcuc
  P  T  K  F  P  I  P  S  P  N  P  R  D  K  W  C  R  L  N  L | SP

  K  P  V  W  G  G  K  C  W  H  G  R  L
 AAGCCTGTGTGGGGGGGAAAGTGCTGGCACGGGAGACTAtaa                   | zcuc
  G  P  A  W  G  G  R  C  -  -  -  -  -                        | SP



  M  D  -  -  -  -  -  A  L  L  K  N  S  V  A  L
 ATGGAC---------------GCCCTGTTGAAGAATTCTGTCGCATTgtgcatgtttcaa| zcuc
  W  E  W  P  W  N  R  K  P  T  K  F  P  I  P  S               | SP

                                              J  L  Y  Q  D  R
 aggcaacaactatagcacgagaaatggtggcaaaattgcagGAGCTTATACCAAGACAGA| zcuc     Not a signal peptide
                                              P  N  P  R  D  K | SP

  W  C  K  L  P  Y  K  P  V  W  G  G  K  C  W  H  G  R  L
 TGGTGCAAATTGCCATACAAGCCTGTGTGGGGGGGAAAGTGCTGGCACGGGAGACTAtaa| zcuc
  W  C  R  L  N  L  G  P  A  W  G  G  R  C  -  -  -  -  -    | SP
```

Figure S12. Gene structure prediction of the potential *SP* homolog in the melon fly, *Zeugodacus cucurbitae*.

A

```
     M  R  A  S  I  M  I  L  M  H  I  K  V  I  I  I  N  M  I  N
310 ATGAGAGCATCGATTATGATTCTTATGCACATCAAAGTCATCATCATCAACATGATCAAC| pvar
  1  M  K  T  L  A  L  F  L  V  L  -  -  V  C  V  L  G  L  V  Q | SP

     N  J  N
370 AACAGCAACAgcaatattcacacggtgatagtgccactacgacaacaacaacaacaacta| pvar
 19  A  W  E                                                     | SP

                                                G  L  A  G  R  T  T
430 cgccgccaccaccaattagctatcattgccgcggcgaagGTCTTGCCGGTCGCACAACAC| pvar
 22                                             W  P  W  N  R  K  P  | SP

    R  W  J  A  L  P  A  P  S  C  R  G  K  W  L  R  L  T  V  G
490 GTTGGAGCGCATTGCCTGCGCCATCGTGTCGTGGCAAATGGTTGCGTTTGACTGTCGGAC| pvar
 29 T  K  F  P  I  P  S  P  N  P  R  D  K  W  C  R  L  N  L  G  | SP

    P  P  -  -  K  K  C  Q  H  A  Q  F  I  F  V
550 CACCG------AAGAAGTGTCAACATGCTCAGTTCATTTTTGTCtaa              | pvar
 49 P  A  W  G  G  R  C  -  -  -  -  -  -  -  -                  | SP
```

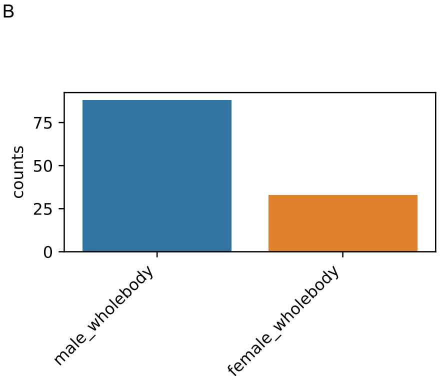


B

>Phortica variegata
MRASIMILMHIKVIIINMINNSNSLAGRTTRWSALPAPSCRGKWLRLTVGPPKKCQHAQFIFV → Signalp

Figure S13. Gene structure prediction of the potential *SP* homolog in the vinegar fly, *Phortica variegata*. A) Sequence alignment of the potential *SP* homolog to *D. melanogaster SP*. The *SP* homolog in *Phortica variegata* also contain two exons and one intron. B) Read counts of the sequence from male and female whole-body RNA-seq data (https://www.ncbi.nlm.nih.gov/bioproject/PRJNA268392). (C) The translation product of the sequence lacks a signal peptide, which is a universal feature of the identified sex peptides.

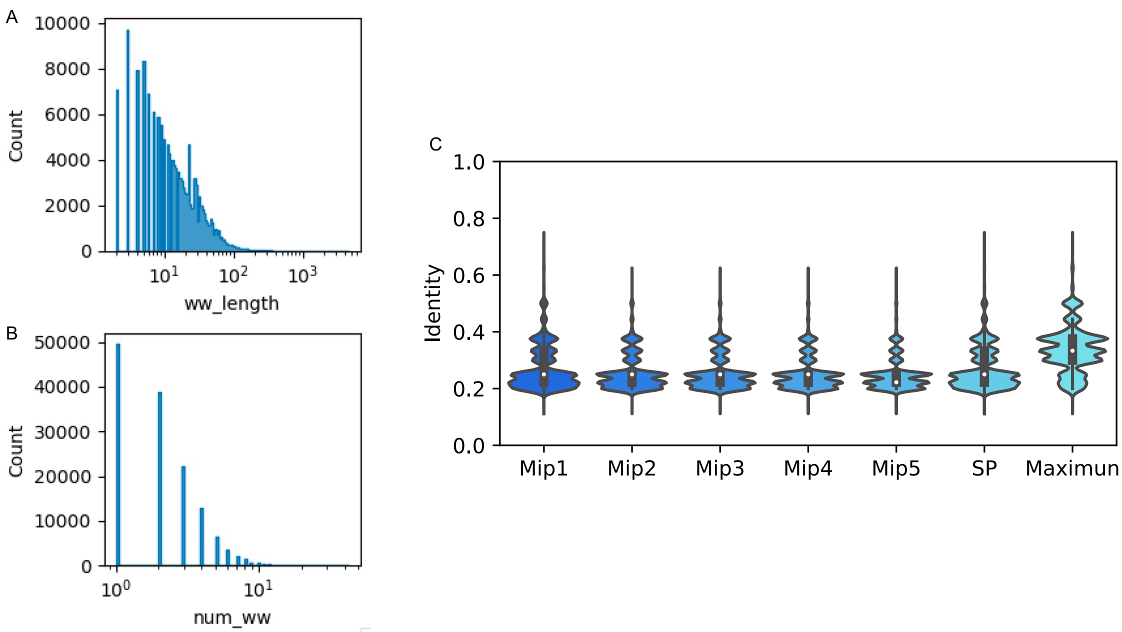


Figure S14. Potential $W(X)_nW$ motif in the *D. melanogaster* genome. (A) Number of intergenic ORFs that contain $W(X)_nW$ motif as the function of the motif length n. (B) Number of intergenic ORFs that contain $W(X)_nW$ motif as the function of the number of motifs inside a single ORF. (C) Sequence identity of the W(X)nW motif to MIP peptides and the $W(X)_nW$ in SP. Maximum sequence identities were shown in the last column.
